# Endocytic trafficking determines cellular tolerance of presynaptic opioid signaling

**DOI:** 10.1101/2022.06.15.496340

**Authors:** Damien Jullié, Camila Benitez, Tracy A. Knight, Mark von Zastrow

## Abstract

Opioid tolerance is well described physiologically but its mechanistic basis remains incompletely understood. An important site of opioid action in vivo is the presynaptic terminal, where opioids inhibit transmitter release. This response characteristically resists desensitization over minutes yet becomes gradually tolerant over hours, and how this is possible remains unknown. Here we delineate a cellular mechanism underlying this longer-term form of opioid tolerance. Our results support a model in which presynaptic tolerance is mediated by a gradual depletion of cognate receptors from the axon surface through iterative rounds of receptor endocytosis and recycling. For the μ-opioid receptor (MOR), we show that the agonist-induced endocytic process which initiates iterative receptor cycling requires GRK2/3-mediated phosphorylation of the receptor’s cytoplasmic tail, and that partial or biased agonist drugs which have reduced ability to drive phosphorylation-dependent endocytosis in terminals produce correspondingly less presynaptic tolerance. We then show that the δ-opioid receptor (DOR) conforms to the same general paradigm except that endocytosis of DOR from the presynapse, in marked contrast to MOR, does not require phosphorylation of the receptor’s cytoplasmic tail. Further, we show that DOR recycles less efficiently than MOR in axons and, consistent with this, that DOR tolerance develops more strongly. Together, these results delineate a cellular basis for the development of presynaptic tolerance to opioids and describe a methodology useful for investigating presynaptic neuromodulation more broadly.

## Introduction

The development of physiological tolerance to opioid agonists provides a fascinating example of neurobehavioral plasticity initiated through the activation of specific G protein-coupled receptors (GPCRs). It is also clinically significant because tolerance limits the therapeutic utility of opioid drugs. While opioid agonists are highly effective in the acute management of pain, maintaining analgesic efficacy under conditions of prolonged or repeated administration tends to require escalating doses. Tolerance develops to other physiological effects of opioids as well, such as the suppression of central ventilatory drive underlying the clinical phenomenon of opioid-induced respiratory depression (OIRD), but typically over a longer period of time (Athanasos et al., 2006; Hayhurst and Durieux, 2016; Paronis and Woods, 1997). This kinetic ‘mismatch’ in the development of tolerance to various opioid-induced effects narrows the therapeutic window for analgesia and is arguably a root cause of the present epidemic of opioid drug-related deaths. Therefore, an important goal of fundamental research is to more fully understand how opioids produce physiological adaptations which develop with widely different rates.

Part of the answer undoubtedly lies in the complexity of in vivo opioid physiology and that opioids impact neural function at multiple levels, from molecular mechanisms in discrete receptor-expressing neurons to adaptations propagating through synaptic networks and neuronal circuits (Cahill et al., 2016; Corder et al., 2018). Even for mechanisms resolved in individual cells, however, it has long been recognized that adaptations can develop at different rates (Chavkin and Goldstein, 1982; Law et al., 1982; Sharma et al., 1975). Accordingly, one plausible approach toward elucidating kinetic differences among opioid-induced neuroadaptations is to focus on molecular mechanisms in individual receptor-expressing neurons that produce physiological effects over a wide range of timescales.

Agonist-induced phosphorylation of receptors is one such mechanism. In particular, phosphorylation of the μ-type opioid receptor (MOP-R or MOR) cytoplasmic tail by GPCR kinases (GRKs) mediates desensitization of MOR-mediated control of potassium channels, a response determining the postsynaptic excitability of neurons (Arttamangkul et al., 2018; Williams et al., 2013). This desensitization process characteristically develops over minutes (Blanchet and Lüscher, 2002; Harris and Williams, 1991; Lowe and Bailey, 2015; Williams et al., 2013), consistent with the time course of MOR phosphorylation and subsequent phosphorylation-dependent endocytosis of MOR in neurons (Arttamangkul et al., 2008; Just et al., 2013). However, phosphorylation of the MOR tail, and on the same Ser/Thr residues required for rapid desensitization, has been clearly shown to attenuate physiological opioid actions after chronic as well as acute administration (Kliewer et al., 2019). Might there be an additional cellular locus at which phosphorylation of the MOR tail drives the development of opioid tolerance over a longer time period?

A possible locus is the presynaptic terminal, where a key physiological action of opioids is to inhibit vesicular neurotransmitter release. Presynaptic inhibition is characteristically resistant to desensitization when assessed over minutes (Blanchet and Lüscher, 2002; Fyfe et al., 2010; Jullié et al., 2020; Lowe and Bailey, 2015; Pennock et al., 2012; Rhim et al., 1993) but has been shown to develop tolerance after prolonged opioid exposure (Fyfe et al., 2010; Lowe and Bailey, 2015). Nevertheless, MOR was recently shown to undergo phosphorylation-dependent endocytosis in presynaptic terminals within minutes (Jullié et al., 2020). Together, these observations suggest the possibility that phosphorylation of the MOR cytoplasmic tail, despite not producing a rapid desensitization of opioid signaling at the presynapse, drives the development of this slower form of presynaptic opioid tolerance.

Here we describe an experimental approach to explicitly test this hypothesis. We delineate a primary culture system enabling the direct measurement of presynaptic tolerance and show that phosphorylation of the MOR cytoplasmic tail is indeed required for this adaptation. We propose a simple cellular mechanism, based on iterative receptor recycling, that is sufficient to explain how rapid phosphorylation of MOR produces presynaptic tolerance over an extended time scale. We then show that a similar model applies to the development of tolerance to presynaptic inhibition by the homologous δ-type opioid receptor (DOP-R or DOR) except that, remarkably, DOR endocytosis in axons does not require phosphorylation of the receptor cytoplasmic tail. Our results provide fundamental insight into the mechanistic basis for opioid-induced neuroadaptations developing over distinct timescales, and describe a methodology that we anticipate will facilitate the study of presynaptic neuromodulation more generally.

## Results

### Presynaptic tolerance to opioids is associated with a loss of surface opioid receptors in the axon

We assayed opioid-induced presynaptic inhibition by adapting a widely used pHluorin-based unquenching assay (Sankaranarayanan et al., 2000) to monitor opioid effects on presynaptic activity in cultured neurons. In this assay, neurons were expressing opioid receptors together with VAMP2-SEP, imaged using a widefield microscope, and were electrically stimulated to induce synaptic vesicle exocytosis. The super-ecliptic pHluorin (SEP) is a pH sensitive GFP whose fluorescence increases as the synaptic vesicle protein VAMP2-SEP relocalizes from acidic synaptic vesicles to the terminal plasma membrane. This fluorescence increase provides a readout for presynaptic activity, which is typically lower when neurons are perfused with agonist for opioid receptors, reflecting opioid mediated presynaptic inhibition. The basic hardware configuration is summarized in Fig 1A, and details of a lab-built apparatus that is economical and produces reliable data are included in Supplementary Methods. We devised an automated analysis pipeline to achieve unbiased identification of synaptic responses (see Supplementary material for further explanation and specific code modules). Responses were measured at individual synapses and then averaged to provide an aggregate readout. Fig 1B shows an example of a recording from the analysis and illustrates how the degree of presynaptic inhibition was defined. We believe this simple assay presents a number of advantages over more complex models to monitor presynaptic inhibition by GPCRs. First, optical measurement of presynaptic activity provides a direct and reliable readout of the degree of inhibition that is independent of compounded postsynaptic effects. Second, the cultured neuron system is highly amenable to genetic and pharmacological manipulations. Third, the hardware and analysis pipeline are simple and largely open-source, facilitating rapid and economical deployment.

**Figure1 :**
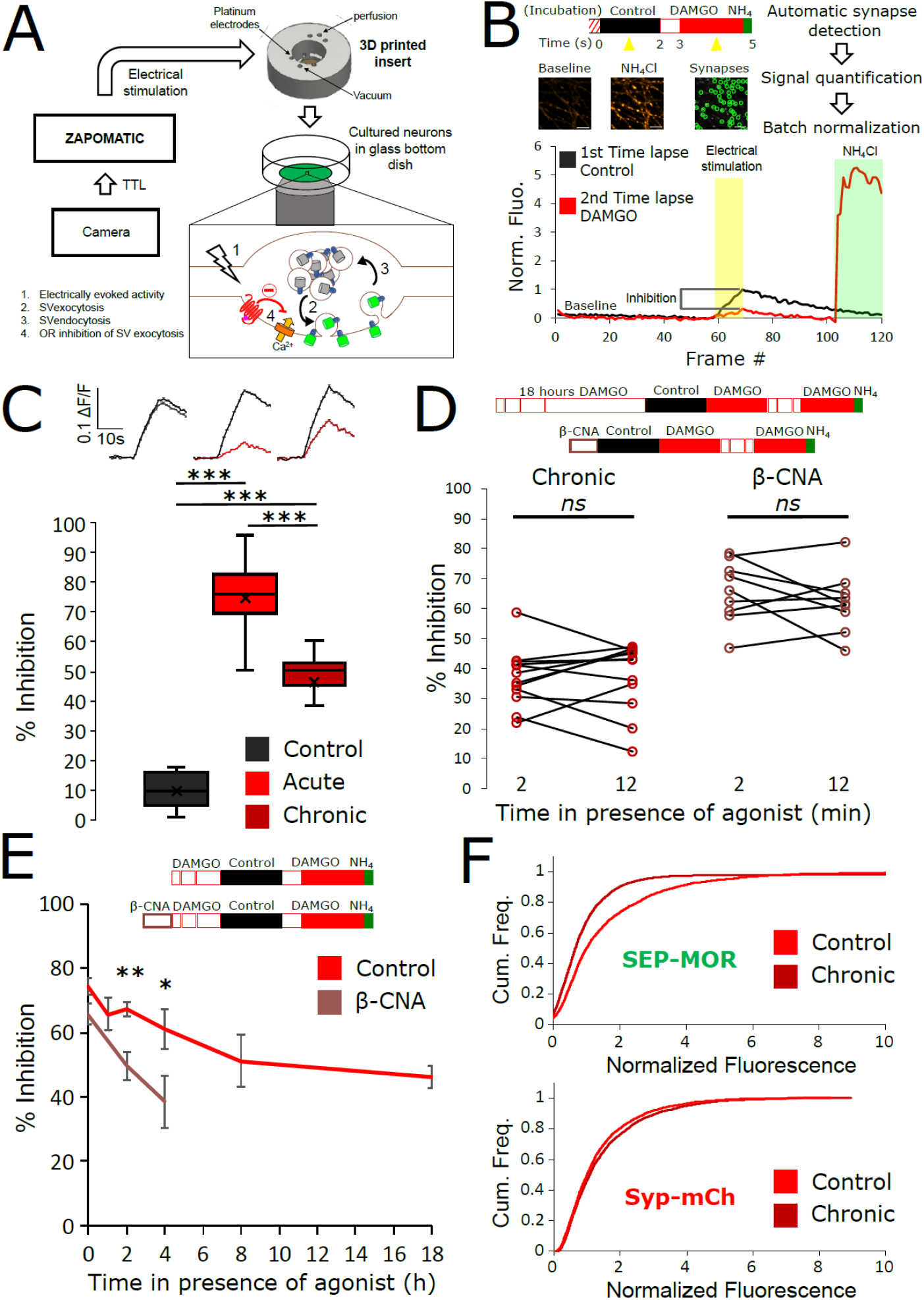
Loss of MOR mediated presynaptic inhibition under chronic activation conditions is paralleled by a reduction in surface receptor number in axons. **A.** Schematic of the experimental setup, with highlighted open-source hardware used for electrical stimulation in synchronicity with image acquisition (zapomatic) and perfusion of solution onto cultured primary cultured neurons (3D printed insert) transfected with opioid receptors and VAMP2-SEP. The enlarged diagram depicts the biological process of electrically stimulated synaptic vesicle recycling (1) monitored with widefield fluorescence microscopy. Calcium entry triggers the exocytosis of VAMP2-SEP containing synaptic vesicles (2) and an increase in fluorescence intensity (green), which returns to baseline after recapture of VAMP2-SEP by endocytosis (3) and quenching of the fluorescence (gray). Active opioid receptor (red) inhibits calcium entry and subsequent exocytosis of synaptic vesicles (4). **B.** Description of the experiment design and automated analysis pipeline. For measurement of acute inhibition, neurons are directly placed in imaging solution on the imaging system. For measurement of inhibition after chronic treatment, neurons are pre-treated with agonist for 18 hours (unless specified otherwise). A first time lapse (120 frames, 1Hz) is acquired in control imaging solution (black box, black curve) and neurons are electrically stimulated at 10Hz for 10s 1 minute into the time lapse. One minute after perfusion of a solution containing DAMGO 10μM (open red box), a second time lapse is acquired (red box, red curve) with the same electrical stimulation and imaging paradigm, and for the last 20 frames the solution is exchanged for ammonium chloride (NH_4_Cl). A differential image NH_4_Cl – baseline is used to automatically detect putative synapses, representatives images are shown. Signal is quantified over multiple tens of putative synapses and used to validate real synapses. Normalized data are pooled for the same condition. For each acquisition, we obtain curves as depicted after normalization by the maximum amplitude of the control condition (n=508 synapses for this acquisition). Note the difference in maximal amplitude in presence of DAMGO compared to control, which reflects inhibition of synaptic vesicle exocytosis by opioid receptors. Scale bar is 10μm. **C.** Upper panel shows average fluorescence curves normalized over NH_4_Cl (ΔF/F) for all synapses, lower panel displays percentage whisker plots of inhibition of SV exocytosis for each acquisition (4 quartiles + mean marker “**X**”). Inhibition of SV exocytosis, compared to control baseline as explained in B, for cells perfused with control solution (Control, inset n = 1603 synapses, n = 6 acquisitions), cells perfused with DAMGO 10μM (Acute, inset n = 3236 synapses, n = 20 acquisitions), and cells pretreated with DAMGO 10μM for 18 hours and perfused with DAMGO (Chronic, inset n = 2025 synapses, n = 13 acquisitions). **D.** To assess rapid desensitization, 3 acquisitions were performed as depicted in the inset. Cells were perfused for 10 minutes in the continuous presence of DAMGO 10μM between stimulations. Paired measurements are shown for cells pretreated with DAMGO 10uM for 18 hours before acquisition (chronic, n = 12 acquisitions), and cells pretreated with β-CNA (50nM for 5 min) before acquisition (β-CNA, n = 9 acquisitions). **E.** Time course of MOR mediated presynaptic inhibition for cells incubated with DAMGO 10μM (n= 20/11/10/11/9/13 acquisitions for 0/1/2/4/8/18 hours, respectively. Time zero and 18 hours replotted from C) or cells pretreated with β-CNA (50nM for 5 min) before incubation with DAMGO (n= 9/7/6 acquisitions for 0/2/4 hours, respectively. Time zero replotted from t=2 minutes in B). **F.** Cumulative frequency curves of the normalized fluorescence at individual synapses for SEP-MOR signal (upper panel) and synaptophysin-mCherry (syp-mCh) for naïve cells (n=3520 synapses) or cells pretreated with DAMGO 10μM for 18 hours (n = 3053 synapses). Note the left shift for SEP-MOR fluorescence after pretreatment indicating a loss of surface receptors. Syp-mCh fluorescence remains similar between conditions, reflecting appropriate sampling of the expression levels of recombinant fluorescent protein among synapses.*, **, *** represent p<0.05, 0.01, 0.001, respectively.

To examine the effect of prolonged opioid exposure in this system, we measured the presynaptic inhibition mediated by [D-Ala^2^, *N*-MePhe^4^, Gly-ol]-enkephalin (DAMGO), a peptide full agonist of MOR. We compared inhibition of the electrically-evoked pHluorin response observed in opioid-naïve neurons (we define this as the acute condition) to that observed in neurons pre-exposed to DAMGO for 18 hours (we define this as the chronic condition). Significant inhibition was observed in both conditions (unpaired t-test compared to control, acute p < 1 e^-5^, chronic p < 1 e^-5^), but the degree of inhibition was reduced in the chronic condition (Fig 1C, unpaired t-test p < 1 e^-5^). These results indicate that prolonged agonist exposure promotes tolerance to presynaptic inhibition by opioids.

Presynaptic inhibition by opioids is well known to be resistant to rapid desensitization processes which attenuate signaling typically over several minutes (Blanchet and Lüscher, 2002; Fyfe et al., 2010; Jullié et al., 2020; Lowe and Bailey, 2015; Pennock et al., 2012; Rhim et al., 1993), suggesting that presynaptic tolerance represents a distinct regulatory process. In addition, after the induction of opioid tolerance in vivo, presynaptic inhibition remains resistant to rapid desensitization while desensitization of the postsynaptic response is enhanced (Arttamangkul et al., 2018; Fyfe et al., 2010). In our in vitro system, we did not detect any evidence for desensitization of the DAMGO response over a 10 minute interval of sequential stimulation after the induction of tolerance (Fig 1D, left. Mean inhibition at 2 minutes 36.98 ± 2.80%, mean inhibition at 12 minutes 37.44 ± 3.34%, paired t-test, p = 0.87). Rather, time course analysis revealed development of tolerance gradually over multiple hours (Fig 1E). This extended time course is reminiscent of the process of receptor downregulation, described previously in other systems and associated with a depletion of the total receptor reserve (Chavkin and Goldstein, 1982; Christie, 2008; Law et al., 1984). Supporting the hypothesis that presynaptic tolerance involves a similar process, we found that reducing receptor reserve using the irreversible antagonist β-Chlornaltrexamine (β-CNA) accelerated the development of presynaptic tolerance (Fig 1E, unpaired t-test, p = 0.075, 0.002, 0.047 for acute, 2 and 4 hours, respectively). This effect was quite sensitive, with significant acceleration evident even under alkylation conditions that have only a small impact on the maximal opioid response and which produce no detectable rapid desensitization of the response (Fig 1D, right, mean inhibition at 2 minutes 65.79 ± 3.41%, mean inhibition at 12 minutes 62.23 ± 3.39%, paired t-test, p = 0.35). To directly test if long term agonist exposure induces a depletion of surface receptors in axons, we imaged SEP-tagged MOR and quantified the fluorescence over thousands of synapses over multiple microscopic fields (Fig 1F). This analysis revealed that prolonged DAMGO exposure indeed reduces the presynaptic surface MOR pool (Fig 1F, MOR no DAMGO mean normalized fluorescence 1.60, 95% confidence interval 1.54-1.66, MOR + DAMGO 18 hours mean normalized fluorescence 1.17, 95%CI 1.08-1.27, two samples Kolmogorov-Smirnov test p < 1e^-5^). Together, these results indicate that presynaptic MOR tolerance is a process distinct from rapid desensitization and likely mediated by a net reduction of surface receptors on axons.

### Endocytosis, tolerance and surface receptor depletion require MOR C-tail phosphorylation

We have previously shown that MOR undergoes rapid agonist-induced endocytosis in terminals, and accumulates in endosomes located both in terminals and in the axon shaft that are marked by retromer complex associated with their limiting membrane (Jullié et al., 2020). We verified this by labeling surface MOR in axons and monitoring DAMGO-induced redistribution of surface-labeled MOR to endosomes marked by GFP-tagged VPS29, a core retromer component (Fig 2A, movie S1). Significant and time-dependent accumulation of surface-labeled MOR in retromer-marked endosomes was observed within several minutes after application of DAMGO, providing a readout for rapid endocytosis in axons (Fig 2B).

**Figure2 :**
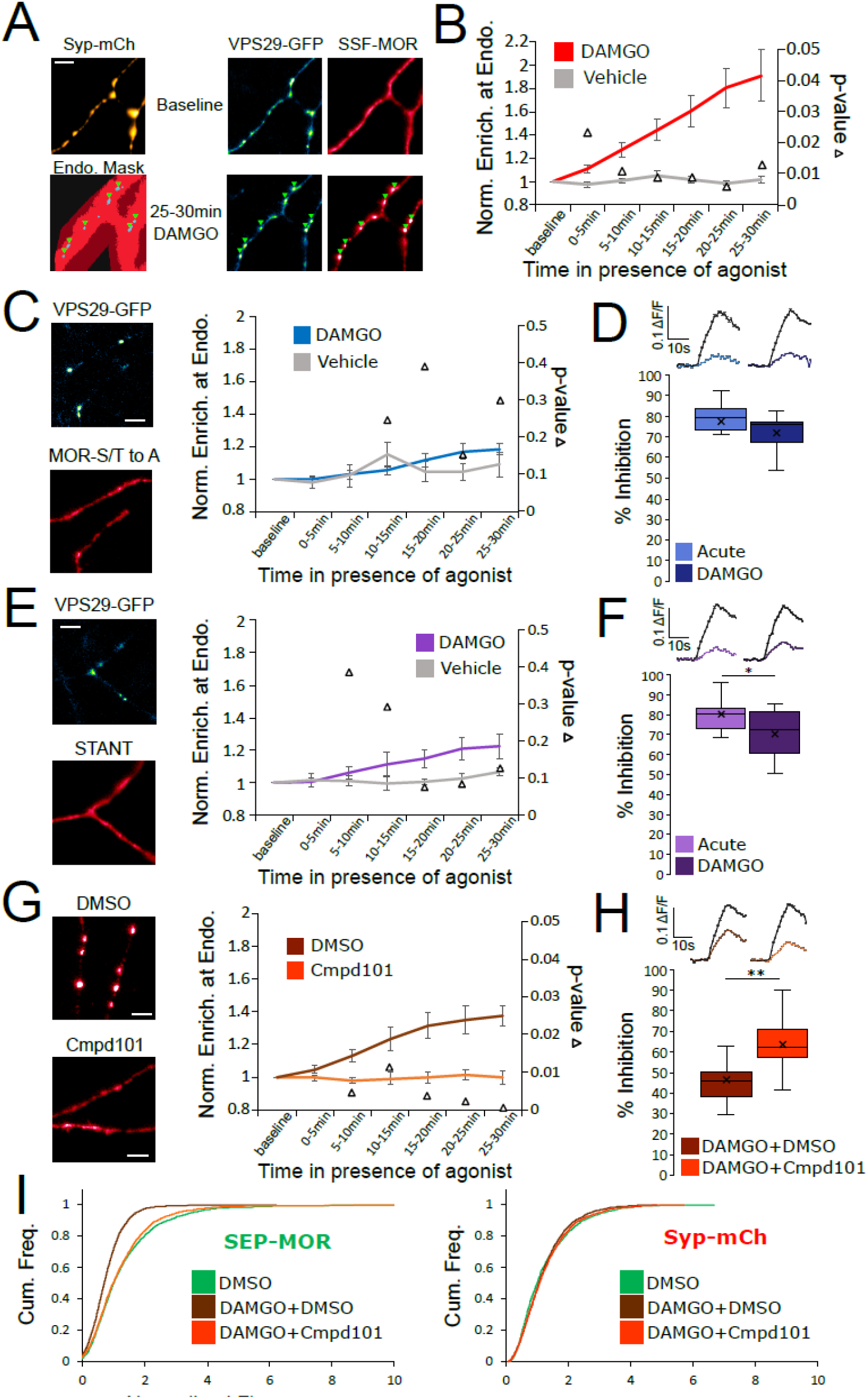
Phosphorylation of MOR is required for endocytosis of receptors, loss of surface receptors upon chronic activation, and the development of presynaptic tolerance. **A.** Representative images of axons of neurons marked with syp-mCh, expressing the endosomal marker VPS29-GFP andFLAG-tagged opioid receptors (SSF-MOR), surface labeled with a primary anti-FLAG antibody conjugated to Alexa 647. Neurons were imaged using oblique illumination at a frequency of 1 frame/minute. Note the uniform distribution of the receptor before agonist addition (baseline) and the punctate distribution overlapping with a segmented mask of the endosomal marker after 25-30 minutes of incubation with DAMGO 10μM. Scale bar is 5μm. See also movie S1. **B.** Quantification of the enrichment of surface labeled SSF-MOR at VPS29-GFP marked structures along axons for cells treated with vehicle (n = 5 acquisitions) or cells treated with DAMGO 10μM (n = 8 acquisitions). Right axis indicates p-values for unpaired t-test between the two conditions. **C.** Same as A,B, for FLAG tagged mutant opioid receptors where all serine and threonine residues of the C-terminal tail have been mutated to alanine (MOR S/T to A), for vehicle (n = 5 acquisitions) or DAMGO 10 μM (n = 7 acquisitions) treated cells. Note the diffused distribution of surface labeled MOR-S/T to A after 25-30min of incubation with DAMGO 10μM. **D.** Quantification of presynaptic inhibition mediated by the MOR S/T to A mutant acutely (inset n = 717 synapses, n = 7 acquisitions) or after 18 hours of incubation with DAMGO 10μM (inset n = 2360 synapses, n = 11 acquisitions). **E.** Same as A,B, for FLAG tagged mutant opioid receptors where serine and threonine residues of STANT motif on the C-terminal tail have been mutated to alanine (STANT), for vehicle (n = 5 acquisitions) or DAMGO 10μM treated cells (n = 6 acquisitions). Note the diffused distribution of surface labeled STANT after 25-30min of incubation with DAMGO 10μM. **F.** Quantification of presynaptic inhibition mediated by the STANT MOR mutant acutely (inset n = 1882 synapses, n = 11 acquisitions) or after 18 hours of incubation with DAMGO 10μM (inset n = 2605 synapses, n = 14 acquisitions). **G.** Same as A,B, for SSF-MOR in neurons treated with Cmpd101 30μM (n=8 acquisitions) or DMSO control (n=6 acquisitions) and incubated with DAMGO 10μM. Note the difference in distribution between the two conditions after 25-30min of incubation with DAMGO. **H.** Quantification of presynaptic inhibition mediated by wild type MOR in cells incubated with Cmpd101 30μM (inset n = 2157 synapses, n = 15 acquisitions) or DMSO control (inset n = 2346 synapses, n = 17 acquisitions) together with DAMGO 10μM for 18 hours. **I.** Cumulative frequency curves of the normalized fluorescence at individual synapses for SEP-MOR signal (left panel) and syp-mCh for cells incubated with DMSO only (n=2273 synapses), cells pretreated with DMSO + DAMGO 10μM for 18 hours (n = 2209 synapses) and cells treated with Cmpd101 30μM + DAMGO 10μM for 18 hours (n = 2456 synapses). Note that the left shift for SEP-MOR fluorescence after pretreatment with DMSO control + DAMGO is blocked by Cmpd101 while syp-mCh control signal is stable across conditions. *, ** represent p<0.05, 0.01, respectively.

Rapid endocytosis of MOR in axons is known to require phosphorylation of Ser and Thr residues in the MOR cytoplasmic tail (Jullié et al., 2020). Using the same assay, we verified that mutation of all Ser and Thr residues in the MOR tail (MOR S/T to A) abolished rapid internalization (Fig 2C). MOR S/T to A strongly inhibited synaptic vesicle exocytosis acutely but mutation of all phosphorylation sites prevented the development of presynaptic tolerance after chronic treatment with DAMGO (Fig 2D, unpaired t-test p = 0.29). Key residues that regulate phosphorylation dependent endocytosis of MOR in other systems are localized into a cluster within the C-terminal tail called the STANT motif (Arttamangkul et al., 2019; Just et al., 2013; Lau et al., 2011). Consistent with this, mutation of the 3 phosphorylatable residues sites in this motif strongly inhibited endocytosis of presynaptic receptors (Fig 2E). The STANT mutant potently inhibited presynaptic activity acutely and little tolerance was observed after 18 hours of treatment with DAMGO (Fig 2F, unpaired t-test p = 0.033). It is known that the GPCR kinases 2 and 3 (GRK2/3) are key regulators of MOR phosphorylation and endocytosis (Jullié et al., 2020; Leff et al., 2020; Lowe and Bailey, 2015; Møller et al., 2020). Consistent with this, compound 101 (Cmpd101), a pharmacological inhibitor of GRK2/3 activity, blocked wild type MOR accumulation in retromer marked endosomes. Further, Cmpd101 significantly blocked the development of tolerance, verifying that phosphorylation is required for the attenuation of presynaptic MOR signaling under conditions of chronic activation (Fig 2H, unpaired t-test p = 0.0012). Cmpd101 also blocked the loss of surface receptors induced by chronic treatment of neurons with DAMGO (Fig 2I, two samples Kolmogorov-Smirnov test: DMSO only mean normalized fluorescence 1.33, 95%CI 1.28-1.38, compared to DMSO + DAMGO mean normalized fluorescence 0.77, 95%CI 0.74-0.79, p < 1e^-5^. DMSO + DAMGO compared to Cmpd101 + DAMGO mean normalized fluorescence 1.23, 95%CI 1.19-1.28 p < 1e^-5^. DMSO only compared to Cmpd101 + DAMGO p = 0.051). Together, these results suggest that GRK2/3 dependent phosphorylation of the MOR tail, by driving the rapid endocytosis of receptors, initiates the process of presynaptic tolerance by reducing the density of receptors present on the axon surface under conditions of prolonged opioid exposure.

### Insight to differences in the effects of chemically distinct opioid agonist drugs

DAMGO efficiently promotes phosphorylation of the MOR tail, and this is a key determinant of β-arrestin recruitment driving subsequent receptor endocytosis. Non-peptide partial agonists such as morphine are less efficacious than DAMGO for promoting receptor phosphorylation as well as endocytosis (Just et al., 2013; Lau et al., 2011). Morphine has been shown to induce presynaptic tolerance in chronically treated animals (Fyfe et al., 2010) but, to our knowledge, its effect on surface MOR availability on axons has not been tested. We were unable to detect significant rapid internalization of MOR in axons, measured after 30 min of morphine exposure, using our endosomal recruitment assay (Fig 3A). However, significant functional tolerance was detected after prolonged (18 hour) morphine exposure (Fig 3B, unpaired t-test, acute compared to morphine + DMSO p = 0.0018). Morphine induced presynaptic tolerance to a reduced degree relative to DAMGO, but it remained dependent on GRK2/3-mediated phosphorylation because it was blocked by Cmpd101 (Fig 3B, unpaired t-test, morphine + DMSO compared to morphine + Cmpd101 p = 0.0016). Accordingly, and despite morphine not producing detectable rapid internalization in axons, we asked whether chronic exposure to morphine is also associated with a phosphorylation-dependent reduction of the overall density of MOR on the axon surface. To test this, we imaged SEP-MOR at synapses after chronic treatment with morphine + Cmpd101 or morphine + DMSO. We found that morphine + DMSO vehicle significantly reduced the amount of receptors at the surface of axons (morphine + DMSO mean normalized fluorescence 1.01, 95%CI 0.96-1.06, compared to DMSO only, two samples Kolmogorov-Smirnov test p < 1e^-5^), but this effect was not as pronounced as when neurons were treated with DAMGO + DMSO (Fig 3C, two samples Kolmogorov-Smirnov test p < 1e^-5^). Furthermore, the morphine-induced reduction of surface receptor number was blocked by Cmpd101 (Fig 3C, morphine + Cmpd101 mean normalized fluorescence 1.23, 95%CI 1.18-1.27, compared to morphine + DMSO two samples Kolmogorov-Smirnov test p < 1e^-5^). These observations suggest that morphine, despite promoting MOR endocytosis only weakly compared to DAMGO, is indeed able to produce presynaptic tolerance after chronic exposure through a similar phosphorylation-dependent mechanism.

**Figure3 :**
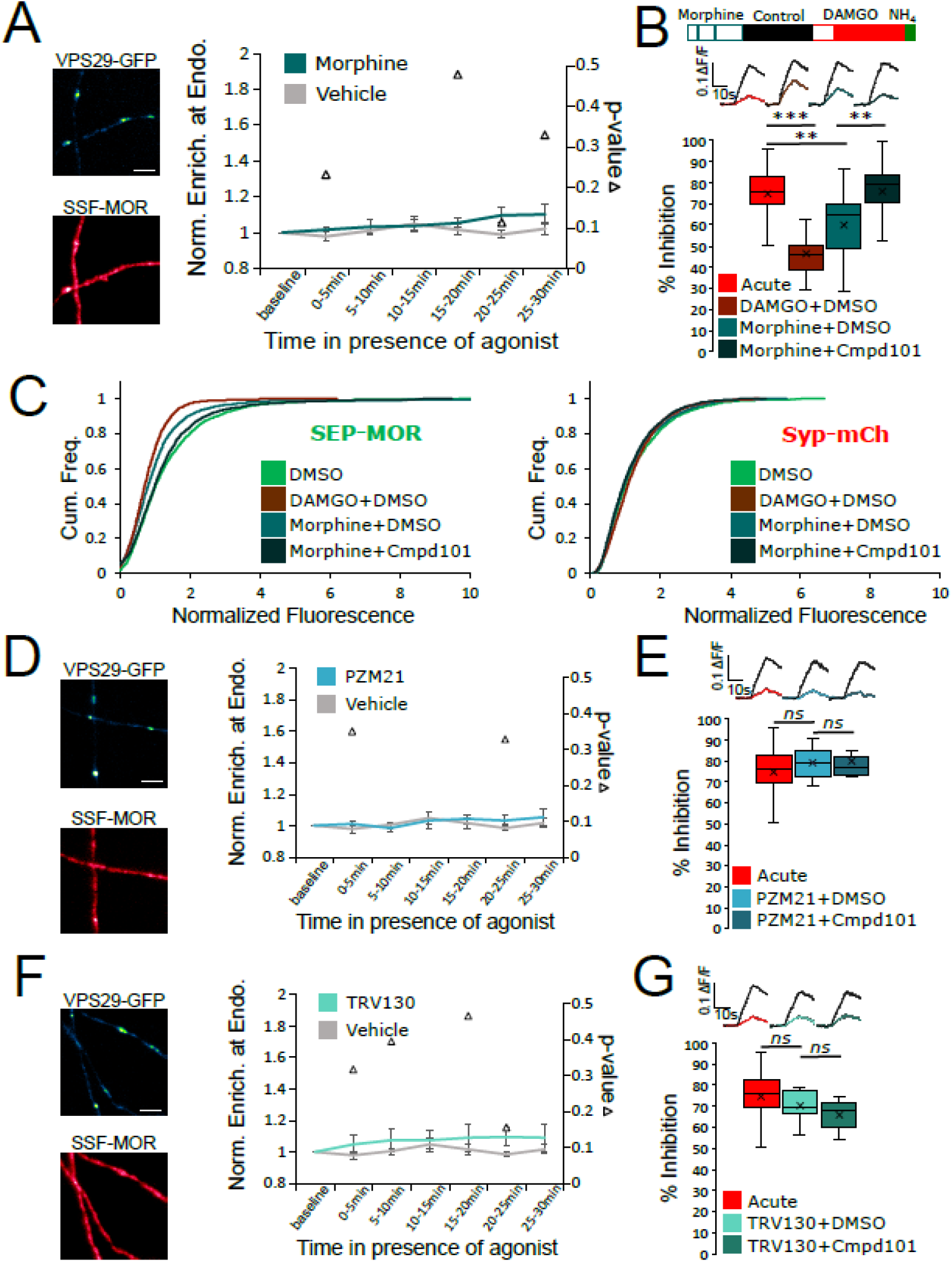
Partial and biased MOR agonists fail to elicit tolerance to the same degree as full agonists peptides. **A.** Same experimental setup as in figure 2, except the agonist used was morphine 10μM (n = 8 acquisitions), control replotted from Fig. 2B. Inset show images of VPS29-GFP and surface labeled SSF-MOR after 25-30min of incubation with morphine 10μM. **B.** DAMGO induced MOR inhibition exhibits tolerance after incubation with morphine 10μM + DMSO vehicle (inset n = 3400 synapses, n = 21 acquisitions) and tolerance is blocked by incubation of morphine 10μM together with Cmpd101 30μM (inset n = 2441 synapses, n = 19 acquisitions). Acute condition replotted from Fig 1C, DMSO + DAMGO condition replotted from Fig 2H. **C.** Morphine 10μM + DMSO (n = 2076 synapses) induces a loss of surface SEP-MOR in axons after 18 hours of incubation compared to DMSO only control (replotted from Fig 2I). The loss is less pronounced than when induced by incubation by DAMGO 10μM + DMSO (replotted from Fig 2I) for 18 hours, and is blocked by incubation of DAMGO 10μM together with Cmpd101 30μM (n = 2872 synapses). Syp-mCh signal is similar across conditions. **D.** Same as Fig 3A except cells were stimulated with PZM21 10μM (n = 5 acquisitions). Note the diffuse distribution of surface labeled SSF-MOR after 25-30min of incubation with PZM21 10μM. **E.** Same as for Fig 3B for cells incubated for 18 hours with PZM21 10μM together with Cmpd101 30μM for 18 hours (inset n = 883 synapses, n = 8 acquisitions) or DMSO vehicle (inset n = 1089 synapses, n = 8 acquisitions). Acute condition replotted from figure 1C. **F.** Same as Fig 3A except cells were stimulated with TRV130 10μM (n = 5 acquisitions). Note the diffuse distribution of surface labeled SSF-MOR after 25-30min of incubation with TRV130 10μM. **G.** Same as for Fig 3B for cells incubated for 18 hours with TRV130 10μM together with Cmpd101 30μM for 18 hours (inset n = 1076 synapses, n = 7 acquisitions) or DMSO vehicle (inset n = 1280 synapses, n = 8 acquisitions). Acute condition replotted from figure 1C. Scale bars are 5μm. **, *** represent p<0.01, 0.001, respectively.

G protein-biased agonists are thought to stimulate MOR internalization even less strongly than morphine. We therefore tested two such compounds, PZM21 and TRV130 (DeWire et al., 2013; Manglik et al., 2016). We could not detect any significant internalization induced by bath application of PZM21, nor tolerance to DAMGO mediated presynaptic inhibition after 18 hours of incubation with the biased agonist (Fig 3D, E, unpaired t-test, acute compared to PZM21 + DMSO p = 0.37, PZM21 + DMSO compared to PZM21 + Cmpd101 p = 0.30). Similarly, TRV130 failed to produce significant MOR internalization or tolerance (Fig 3F, G, unpaired t-test, acute compared to TRV130 + DMSO p = 0.37, TRV130 + DMSO compared to TRV130 + Cmpd101 p = 0.091). Together, these results establish a strong correlation between the known endocytosis efficacy of chemically diverse MOR agonists and the observed degree of tolerance.

### DOR exhibits a higher degree of tolerance than MOR and indicates that tolerance is an homologous process

MOR is not the only receptor mediating presynaptic neuromodulation by opioids. DOR is another well-known example that mediates Gi-coupled inhibition of neurotransmitter release in response to opioids (Bardoni et al., 2014; He et al., 2021; Piskorowski and Chevaleyre, 2013). Physiological tolerance to DOR-mediated effects is well established (DiCello et al., 2019; Pradhan et al., 2010), and agonist-induced internalization of DOR has been clearly demonstrated in the soma and dendrites of neurons (Pradhan et al., 2009; Scherrer et al., 2006). Recent evidence indicates that DOR does not rapidly desensitize at the presynapse (He et al., 2021). However, DOR trafficking in axons has not been studied previously, and it is not known if longer-term tolerance develops to DOR-mediated presynaptic inhibition. We found that, similar to MOR, surface labeled DOR is diffusely distributed in axons of striatal neurons and does not accumulate specifically at synapses marked with syp-mCh (Fig 1A, baseline). After stimulation of DOR with the peptide agonist [D-Ala^2^, D-Leu^5^]-Enkephalin (DADLE), there was a redistribution of surface receptors in punctate structures that colocalized with the retromer marker VPS29-GFP (Fig 1A,B, movie S2). This indicates that, as for MOR, presynaptic DOR undergoes ligand dependent endocytosis and accumulates in a similar population of presynaptic endosomes. Using our optical assay to probe DOR mediated presynaptic inhibition, we found that DOR potently inhibits exocytosis of synaptic vesicles (Fig 4C, D). We could not detect significant attenuation of DOR signaling after 10 minutes of perfusion with agonist, indicating that presynaptic DOR resists acute desensitization (Fig 1C, mean inhibition at 2 minutes 81.90 ± 2.76%, mean inhibition at 12 minutes 82.96 ± 4.55%, paired t-test, p = 0.80). We probed presynaptic DOR tolerance after treatment of neurons for 18 hours with DADLE, and found that after chronic treatment DOR mediated inhibition of synaptic vesicle exocytosis is barely detectable indicating robust tolerance (Fig 4D, unpaired t-test, acute compared to chronic p < 1e^-5^). Accordingly, while resistance to acute desensitization and presynaptic tolerance are properties shared between opioid receptors, DOR-mediated presynaptic inhibition exhibited a greater degree of tolerance after the same period of prolonged agonist exposure (Fig 4D, unpaired t-test, MOR compared to DOR after chronic treatment p = 1.01e^-5^).

**Figure4 :**
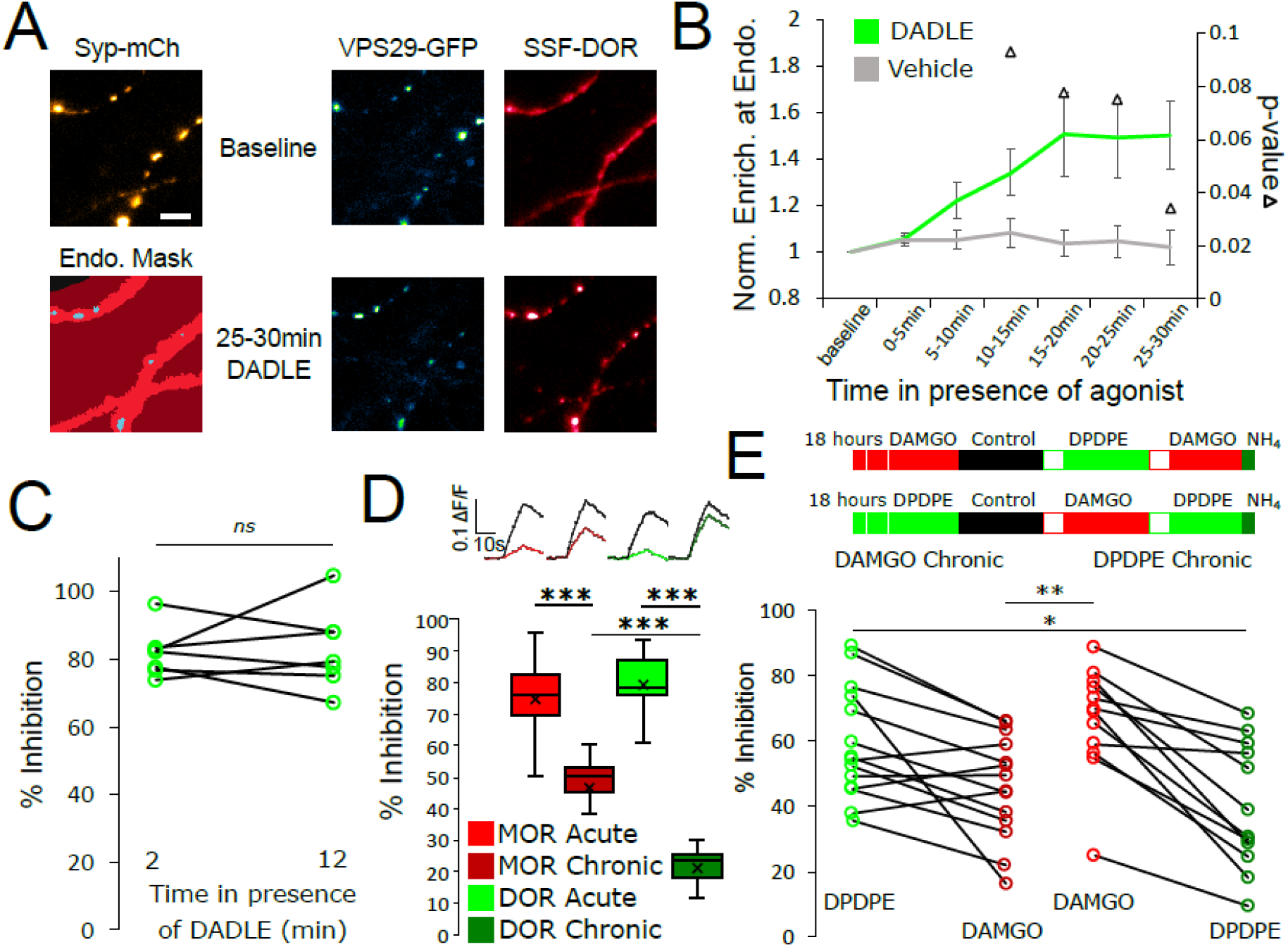
Tolerance is an homologous process conserved between opioid receptors. **A.** Representative images of axons of neurons marked with syp-mCh, expressing the endosomal marker VPS29-GFP and FLAG-tagged DOR (SSF-DOR) surface labeled with a primary anti-FLAG antibody conjugated to alexa647. Imaging was performed as described before. Note the uniform distribution of surface SSF-DOR before agonist addition (baseline) and the punctate distribution overlapping with a segmented mask of the endosomal marker after 25-30 minutes of incubation with DADLE 10μM. Scale bar is 5μm. See also movie S2. **B.** Time course of surface labeled SSF-DOR recruitment at VPS29-GFP marked presynaptic endosomes, as in A. There is a significant increase in colocalization of SSF-DOR with the retromer marker after addition of DADLE 10μM (n = 11 acquisitions) compared to the vehicle control (n = 7 acquisitions). **C.** Inhibition of electrically evoked exocytosis of synaptic vesicles by DOR is sustained over 10 minutes in presence of agonist. Desensitization of presynaptic DOR was assessed using a similar protocol as in Fig 1D. Neurons expressing SSF-DOR and VAMP2-SEP were electrically stimulated to induce SV exocytosis in control solution. Cells were then perfused with a solution containing DADLE 10μM and inhibition of the fluorescence increase was quantified to reflect DOR mediated presynaptic inhibition. After 10 more minutes of perfusion with DADLE, cells were stimulated again to estimate the degree of acute desensitization (n = 7 acquisitions). **D.** Quantification of the presynaptic inhibition mediated by MOR in acute and chronic conditions (replotted from Fig 1C) compared to DOR, acutely (inset n = 1482 synapses, n = 10 acquisitions) or after 18 hours of treatment with DADLE 10μM (inset n = 2529 synapses, n = 11 acquisitions). **E.** Assessment of cross-tolerance using optical measurement of presynaptic inhibition. Inset describes experimental setup. Neurons expressing VAMP2-SEP together with SSF-DOR and SSF-MOR were incubated for 18 hours with either DAMGO 10μM (n = 14 acquisitions) or DPDPE 10μM (n = 12 acquisitions). Cells treated chronically with DAMGO were electrically stimulated while imaged in control solution, then 2 minutes after perfusion with 10uM DPDPE, then 2 minutes after exchange for a solution containing 10μM DAMGO. Cells treated chronically with DPDPE were submitted to the same protocol, except that the order of solution perfusion was reversed (DAMGO first, DPDPE second). *, **, *** represent p<0.05, 0.01, 0.001, respectively.

DOR is co-expressed with MOR in some neurons, and it has been proposed that such co-expression can underlie functional cross-talk and cross-regulation between these distinct opioid receptor types (He et al., 2021; Wang et al., 2018). This motivated us to ask if presynaptic tolerance elicited by chronic agonist exposure is receptor-specific or if chronic activation of one receptor type promotes the development of tolerance at the other. Neurons co expressing MOR and DOR were probed in our optical assay after the chronic treatment with either DAMGO or [D-Pen^2,5^]-Enkephalin (DPDPE), a full agonist peptide that exhibits much higher specificity for DOR than DADLE. We find that chronic MOR activation preserves DOR potent presynaptic inhibition while developing tolerance at MOR (Mean MOR inhibition after DAMGO chronic 45.85 ± 4.19%, mean MOR inhibition after DPDPE chronic 66.09 ± 4.80%, unpaired t-test p = 0.0039). Conversely, chronic treatment of neurons with DPDPE did not induce presynaptic MOR tolerance but attenuated DOR signaling (Fig 4E, mean DOR inhibition after DAMGO chronic 59.10 ± 4.59%, mean DOR inhibition after DPDPE chronic 39.72 ± 5.58%, unpaired t-test p = 0.012). These results indicate that tolerance is a shared phenomenon between opioid receptors but that the process is homologous and regulated at the receptor level.

### Differences in the degree of presynaptic tolerance between receptor types correlate with differences in surface receptor depletion and recycling rate

As our cellular model links presynaptic tolerance to a reduction in the surface opioid receptors pool, we anticipated from the above results that loss of surface receptors occurs to a greater degree for DOR than MOR. To test this, we imaged surface DOR fluorescence using a N-terminally SEP-tagged construct. We found that 18 hours of incubation with DADLE led to reduction in the number of surface receptors, indicating that DOR tolerance is paralleled by a reduction of the receptor pool in axons (DOR no DADLE mean normalized fluorescence 1.43, 95%CI 1.38-1.48, compared to DOR + 18 hours DADLE mean normalized fluorescence 0.32, 95%CI 0.31-0.34, two samples Kolmogorov-Smirnov test p < 1e^-5^). Also, the loss of receptor induced by chronic treatment was much more pronounced than for SEP-MOR, in agreement with our measurements of presynaptic inhibition after induction of tolerance (Fig 5A, MOR + DAMGO compared to DOR + DADLE, two samples Kolmogorov-Smirnov test p < 1e^-5^). Enhanced down-regulation of surface DOR relative to MOR has been observed previously in non-neural models, and results from a reduced efficiency of DOR to enter the recycling pathway compared to MOR (Tanowitz and von Zastrow, 2003). To test if this is the case in axons, we incubated neurons expressing syp-mCh together with SEP-tagged opioid receptors with agonist for 20 minutes, a time that is sufficient to engage receptors into the endosomal pathway. We then imaged axons at a higher acquisition rate (10Hz) sufficient to resolve individual receptor-containing vesicular fusion events mediating receptor recycling to the axon surface; these appear as bursts of fluorescence due to rapid SEP dequenching upon exposure to the neutral extracellular milieu (Fig 5B). Such insertion events were detected for both SEP-DOR and SEP-MOR, with no significant difference in amplitude (Fig 5C, unpaired t-test, p = 0.93). While recycling events were rare for both receptors in neurons that were not pretreated with agonist, their frequency was significantly higher after agonist treatment (unpaired t-test, MOR p = 0.00023, DOR p = 0.0015), consistent with ligand-induced trafficking and recycling. More importantly, we found that after agonist treatment, the frequency of SEP-DOR surface insertion events was about half of what was observed for SEP-MOR (unpaired t-test, p = 0.0231). Together, these data suggest that while both MOR and DOR undergo ligand-dependent endocytosis in axons, DOR recycles less efficiently than MOR. We propose that, when iterated over the course of 18 hours of agonist treatment, this produces a significant difference in the degree of receptor loss from the axon surface underlying the observed difference in degree of tolerance.

**Figure 5:**
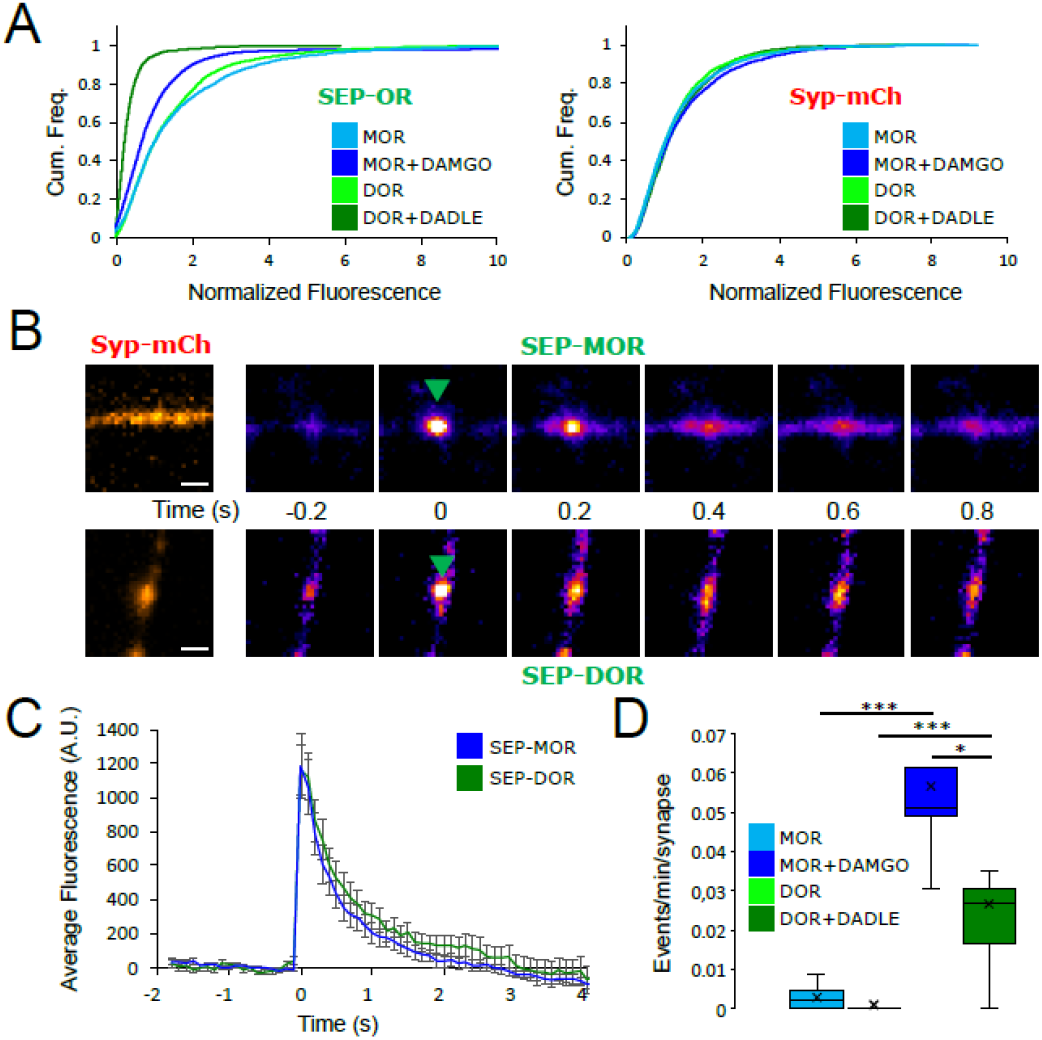
DOR is less efficiently recycled to the plasma membrane compared to MOR. **A.** 18 hours of incubation with DADLE 10μm (n=2934 synapses) induces a marked loss of surface SEP-DOR in axons compared to the untreated control (n=3376 synapses). The loss is more pronounced than what is observed after chronic treatment of SEP-MOR with DAMGO (replotted from Fig 1F). Syp-mCh fluorescence control remains similar across conditions. **B.** Representative examples of surface insertion of SEP-tagged opioid receptors. Neurons were incubated for 20 minutes with either DAMGO 10μM (for SEP-MOR) or DADLE 10μM (for SEP-DOR), and imaged at 10Hz using oblique illumination. Insertion events appear as bursts of fluorescence (green arrow). Scale bar is 1μm. **C.** Average fluorescence intensity profile at the site of insertion for SEP-MOR (n=59 events) and SEP-DOR (n=49 events), for events imaged as described in B. Error bars represent SEM. **D.** Whisker plots of the normalized frequency of surface insertion events for neurons imaged as described in B. Frequency of recycling events was increased for both MOR (n = 5 acquisitions) and DOR (n = 8 acquisitions) compared to the no agonist pretreatment condition (MOR n = 6 acquisitions, DOR n = 8 acquisitions). *, *** represent p<0.05, 0.001, respectively.

### Presynaptic DOR endocytosis, surface receptor depletion and tolerance do not require C-tail phosphorylation

Whereas both MOR and DOR appear to rely on endocytosis-associated loss of axonal receptor for the development of presynaptic tolerance to opioids, the mechanistic basis for its agonist-dependent control appeared remarkably different between the two opioid receptor types. Specifically, while MOR internalization, surface receptor loss, and tolerance clearly require phosphorylation of the receptor’s cytoplasmic tail, this was not the case for DOR. First, the degree of DADLE-induced DOR tolerance observed in the presence of Cmpd101 was indistinguishable from the DMSO vehicle control (Fig 6A, unpaired t-test, p = 0.71). We also observed rapid internalization of DOR in the presence of Cmpd101 (Fig. 6B). These results indicate that DOR tolerance and internalization do not require GRK2/3 activity, in contrast to MOR. Second, mutating all Ser and Thr residues in the DOR C-terminal tail (DOR S/T to A) did not prevent the development of presynaptic tolerance (Fig. 6C, unpaired t-test, p = 0.00036). We also observed significant rapid internalization of DOR S/T to A, as indicated by the mutant receptor undergoing DADLE-induced accumulation at retromer marked endosomes (Fig. 6D, movie S3). These results indicate that DOR tolerance and internalization do not require phosphorylation of the receptor’s cytoplasmic tail, in contrast to MOR. Third, assay of surface receptor fluorescence indicated that the S/T to A mutation does not prevent DADLE-induced reduction of the surface pool of SEP-DOR (Fig. 6E, DOR S/T to A no DADLE mean normalized fluorescence 1.30, 95%CI 1.27-1.32, compared to DOR S/T to A + 18 hours DADLE mean normalized fluorescence 0.45, 95%CI 0.44-0.46, two samples Kolmogorov-Smirnov test p < 1e^-5^). This indicates that agonist-induced reduction of the axonal surface DOR pool does not require phosphorylation of the receptor cytoplasmic tail.

**Figure 6:**
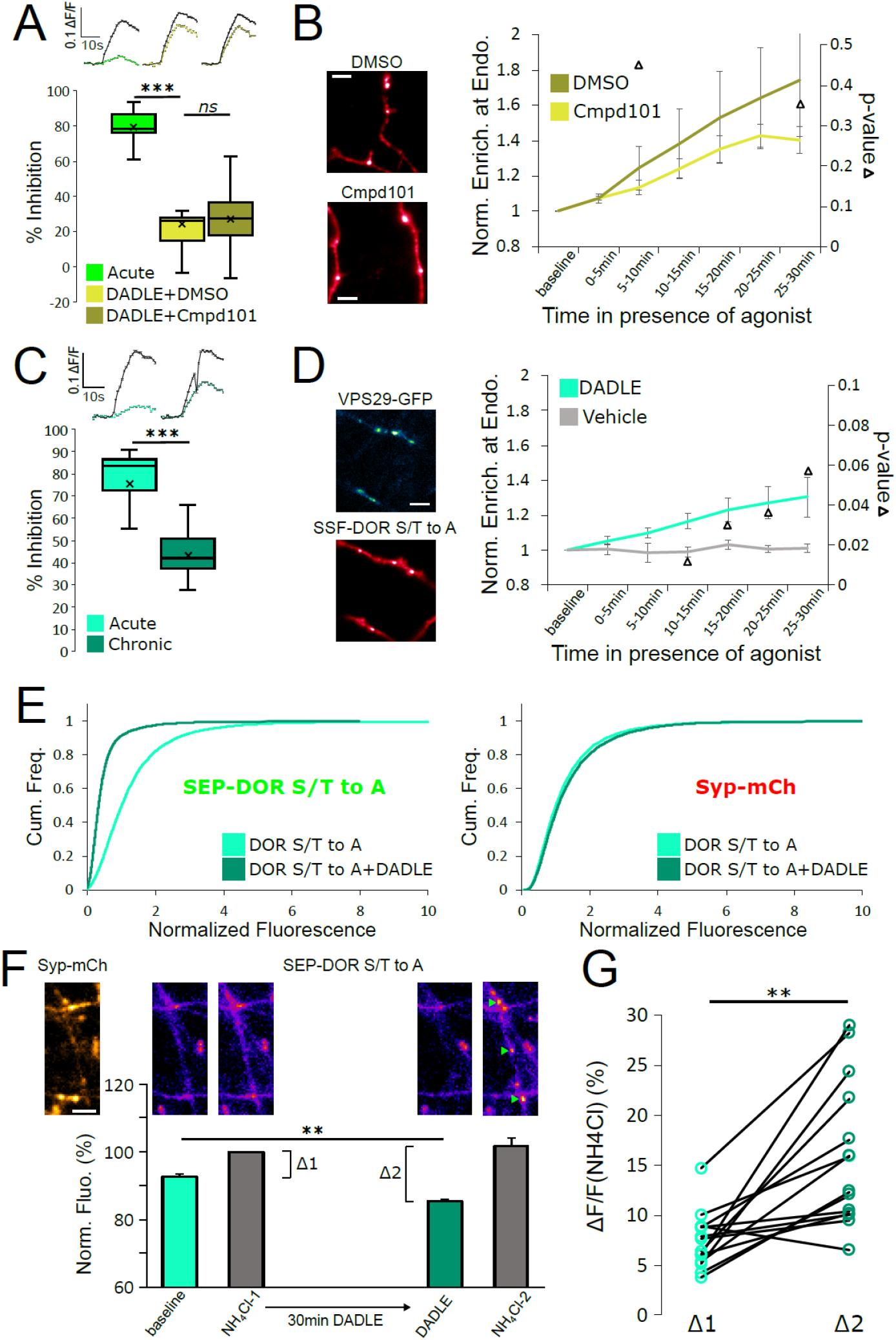
DOR C-terminal tail phosphorylation is not necessary for presynaptic endocytosis, surface receptor depletion or tolerance. **A.** Inhibition of GRK2/3 activity does not block DOR tolerance. Neurons were incubated with Cmpd101 30μM + DADLE 10μM (inset n=2481 synapses, n=15 acquisitions) or DMSO vehicle + DADLE 10μM (inset n=2353 synapses, n=13 acquisitions) for 18 hours, and inhibition of presynaptic activity measured as described previously. Acute condition is reploted from Fig 4D. **B.** Effect of GRK2/3 inhibition on the accumulation of surface labeled SSF-DOR at VPS29-GFP marked endosomes, as previously described. Neurons were incubated with Cmpd101 30μM (n = 7 acquisitions) or DMSO vehicle (n = 7 acquisitions) and DADLE 10μM was added after baseline. Note the punctate distribution for both conditions after 25-30 min of incubation with agonist. Scale bar is 5μm. **C.** DOR S/T to A develops tolerance after chronic activation. Presynaptic inhibition mediated by a phosphorylation-deficient mutant of DOR was assessed in neurons treated acutely (inset n=1673 synapses, n=8 acquisitions) or in neurons pretreated for 18 hours with DADLE (inset n=3017 synapses, n=12 acquisitions). **D.** Endocytosis of DOR S/T to A, as described previously. Neurons expressing the mutant were stimulated with DADLE 10μM (n = 7 acquisitions) or vehicle control (n = 6 acquisitions). Note the punctate distribution overlapping the endosomal marker signal after 25-30 min of incubation with agonist. Scale bar is 5μm. See also movie S3. **E.** Normalized fluorescence of SEP-DOR S/T to A at synapses marked by syp-mCh in naive neurons (n = 7200 synapses) or in neurons pretreated with DADLE 10μM for 18 hours (n = 8904 synapses). Note the left shift in SEP-DOR S/T to A fluorescence after chronic activation while the syp-mCh signal remains stable**. F.** Endocytosis of SEP-DOR S/T to A assessed by pHluorin unquenching. Axons were identified by syp-mCh staining and perfused with imaging solution, and a baseline image acquired. 1min after perfusion of NH_4_Cl solution, another image was taken (NH_4_Cl-1) showing little increase in fluorescence (Δ1). Cells were then perfused for 30min with DADLE 10μM in imaging solution, and another frame acquired (DADLE). One last frame was acquired after 1min of perfusion with NH_4_Cl (NH_4_Cl-2) showing a larger increase in fluorescence (Δ2). Inset shows representative images for each step, green arrows point to fluorescent punctates that represent endosomes. Scale bar is 5μm, n = 14 acquisitions. **G.** Paired measurement of the fluorescence increase induced by NH_4_Cl at baseline (Δ1) or after 30min of incubation with DADLE 10μM (Δ2), as described in E, same dataset. **, *** represent p< 0.01, 0.001, respectively.

Altogether our data suggest that endocytosis is responsible for long term surface receptor depletion for both MOR and DOR. However, the above results indicate that DOR endocytosis does not require phosphorylation of the receptor tail. To confirm this result, we used a different assay that leverages the properties of SEP. Ammonium chloride (NH_4_Cl) can titrate acidic intracellular compartments and reveal SEP fluorescence from internal receptors. In neurons expressing SEP-DOR S/T to A, and in basal conditions, application of NH_4_Cl induced a modest fluorescence increase (Δ1, mean = 7.39 ± 0.75%), suggesting that SEP-DOR S/T to A mostly resides at the surface of the axon. After 30min of DADLE bath application, NH_4_Cl application led to a significantly larger fluorescence increase (Δ2, mean = 16.45 ± 2.27%, paired t-test p = 0.0017), indicating that a proportion of mutant receptors had relocalized from the surface to internal acidic organelles (Fig. 6F,G). These results independently confirm that C-tail phosphorylation is not required for DOR endocytosis in axons.

## Discussion

The present results establish that GRK-mediated phosphorylation of the MOR cytoplasmic tail is required for a form of cellular opioid tolerance at the presynaptic terminal which develops over a significantly longer period of time than the previously elucidated process of rapid desensitization of postsynaptic MOR signaling. Our results support a simple cellular mechanism that is sufficient to explain the slower kinetics with which presynaptic tolerance develops, based on an iterative endocytic trafficking cycle which mediates a progressive depletion of receptors from the axon surface. Accordingly, the present results provide new insight into the cellular and molecular mechanisms underlying the diversity of timescales over which functionally relevant neuroadaptations to opioids develop.

Presynaptic tolerance develops gradually because it represents an integrated effect of repetitive rounds of endocytosis and recycling, in which a small fraction of internalized receptors are not reinserted in each cycle (Fig 7, large arrow). The key event initiating this iterative trafficking cycle is endocytosis of the receptor and, for MOR, this requires phosphorylation of the receptor’s cytoplasmic tail. Supporting this conclusion, blocking MOR phosphorylation in multiple ways also prevents the gradual depletion of surface receptors and the development of functional presynaptic tolerance. Further, agonists which do not drive MOR endocytosis robustly produce less (morphine) or no (PZM21, TRV130) measureable presynaptic tolerance. The idea that tolerance develops as a consequence of gradual depletion of the total receptor pool on the axon surface is also consistent with our finding that β-CNA, a distinct manipulation that reduces total receptor reserve, dramatically accelerates the development of opioid-induced presynaptic tolerance. Thus the present model is supported at multiple levels, and it is sufficient to explain the extended time course of presynaptic tolerance.

**Figure 7:**
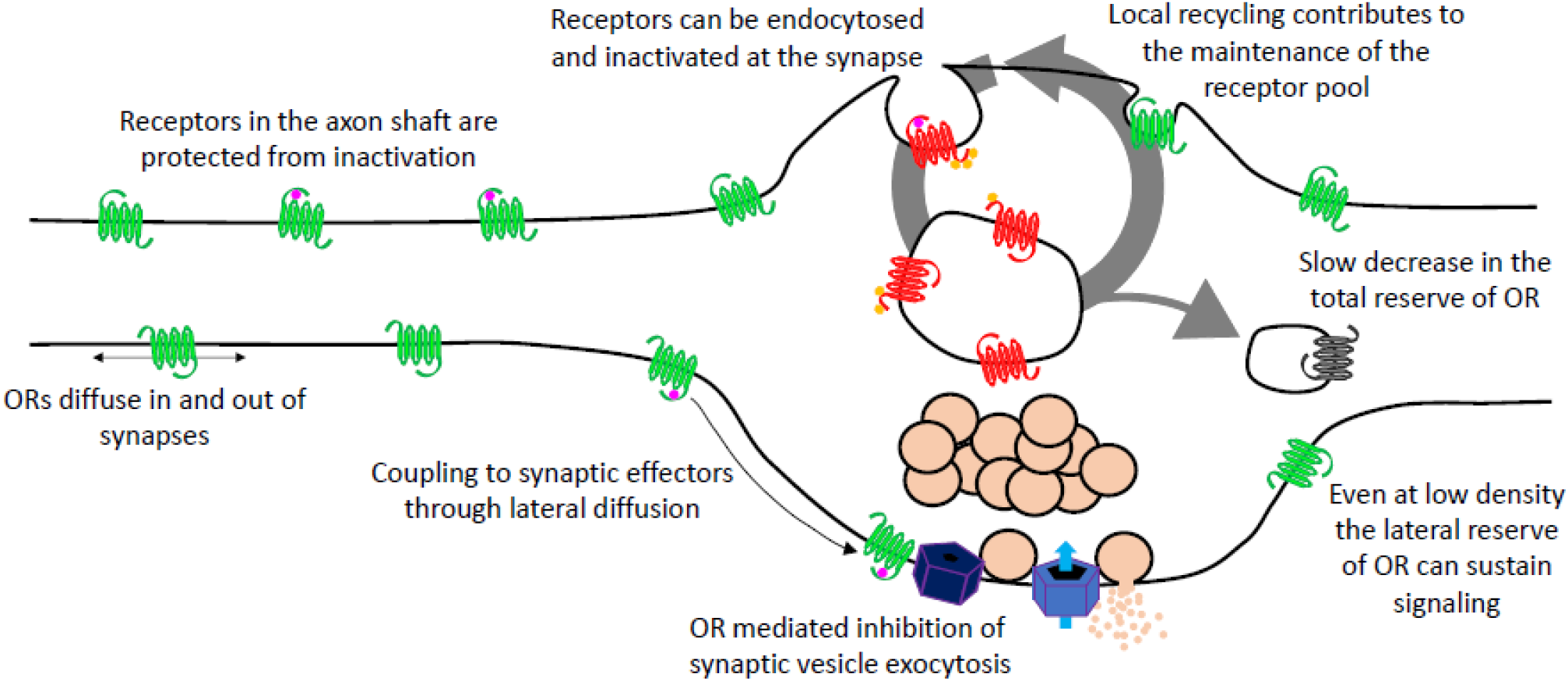
Summary model of opioid receptor signaling and trafficking in axons.

A cellular basis for the ability of presynaptic inhibition by MOR to resist rapid desensitization was proposed previously, grounded in two distinguishing properties of MOR cell biology in axons. First, endocytosis of receptors occurs almost exclusively in synaptic specializations that are sparsely distributed along the extended axon shaft. Second, opioid receptors are laterally mobile over the entire axon surface, with receptors present at synapses able to exchange with the adjacent extrasynaptic pool within seconds (Jullié et al., 2020). Accordingly, receptors present on the axon surface, but outside of synapses, are able to diffuse into synapses at a faster rate than receptors undergo agonist-induced endocytic capture and inactivation within synapses. The extrasynaptic membrane thus provides a ‘lateral reserve’ of receptors enabling opioid signaling at terminals to be maintained even at low overall surface receptor density (Fig 7, small arrows). Presynaptic tolerance appears to develop, instead, through a progressive reduction of the ‘total reserve’ of receptors on the axon surface (Fig 7, large arrow). By elaborating the classical concept of receptor reserve in this manner, and extending it here to DOR, the present results explain how presynaptic inhibition by opioids resists rapid desensitization while becoming tolerant over a longer time period.

Our proposed model for the development of presynaptic tolerance is reminiscent of a general model of GPCR downregulation that was pioneered through the study of DOR in non-neural cells (Law et al., 1982; Williams et al., 2013). Our results are consistent with this model, including the present demonstration that DOR recycles less efficiently than MOR in axons and develops presynaptic tolerance more robustly. A key point of divergence is that, for DOR, none of the events associated with the development of tolerance at the presynapse– beginning with endocytosis of the receptor– requires phosphorylation of the receptor tail. Agonist-induced phosphorylation of the DOR tail has been clearly demonstrated, and many studies support its importance for mediating DOR endocytosis and/or downregulation in other cellular contexts. e.g., (Mann et al., 2020; Whistler et al., 2001). However, we also note that there is evidence indicating that phosphorylation is not absolutely required for DOR endocytosis (Qiu et al., 2007; Zhang et al., 2005). An interesting question for future study is to determine whether or not the presently observed difference in the molecular requirements for MOR and DOR endocytosis is specific to the presynaptic compartment.

In sum, our results inform the fundamental question of how opioids produce neuroadaptive effects that span a wide range of timescales, and they delineate a cellular mechanism for the development of presynaptic tolerance. The present results are limited to two opioid receptor types. However, we note that MOR and DOR belong to the largest GPCR class (family A) and that presynaptic inhibition mediated by the A1 adenosine receptor (also a family A GPCR) resists rapid desensitization yet develops tolerance gradually (Wetherington and Lambert, 2002). Thus we anticipate that the present results, in addition to their specific relevance to opioid drug effects, provide a framework and methodology useful for understanding presynaptic neuromodulation through GPCRs more generally.

## Acknowledgements

We thank Aashish Manglik for providing aliquots of PZM21 and TRV130. We thank Miriam Stoeber for valuable discussions. We thank members of the von Zastrow lab for valuable discussion, suggestions, and technical help throughout this project. Oblique illumination experiments were performed at the Center for Advanced Light Microscopy at UCSF and we thank its expert staff for providing advice and maintaining the equipment. We thank the Center for Advanced Technology at UCSF for providing access to the 3D printer. This study was supported by research grants from the NIH/NIDA (DA010711 and DA012864).

## Author Contributions

D.J. and M.v.Z. built the hardware, conceived the experiments and wrote the manuscript. D.J. performed the experiments and analyzed the data. D.J. and C.B. developed reagents. D.J. and T.K. developed software.

## Material and Methods

### Primary rat striatal neuron cultures

All procedures were performed according to the National Institutes of Health Guide for Care and Use of Laboratory Animals and approved by the University of California San Francisco Institutional Animal Care and Use Committee. Briefly, after euthanasia of the pregnant Sprague-Dawley rat (CO_2_ and bilateral thoracotomy), the brains of embryonic day 18 rats were extracted from the skull. The striatum, including the caudate-putamen and nucleus accumbens, were identified (Banker and Goslin, 1999). The structures were dissected in ice cold Hank’s buffered saline solution Calcium/magnesium/phenol red free (Gibco). Striatum were dissociated in 0.05% trypsin/EDTA (Gibco) for 15 min at 37°C before 2 washes in Dulbecco’s modified Eagle’s medium (DMEM, Gibco) supplemented with 10% fetal bovine serum (University of California, San Francisco, Cell Culture Facility) and 30mM HEPES (Gibco). Neurons were mechanically separated with a flame-polished Pasteur pipette. Nucleofected striatal neurons were transfected using manufacturer’s instructions (Rat Neuron Nucleofector Kit, Lonza) for rat hippocampal neurons before plating. Cells were plated on poly-D-lysine coated 35mm glass bottom dishes (Matek) in DMEM supplemented with 10% fetal bovine serum. Medium was exchanged 1-4 days after plating for phenol red free Neurobasal (Gibco) supplemented with Glutamax 1x (Gibco) and B27 1x (Thermo Fisher). Half of the culture medium was exchanged every week with fresh, equilibrated medium. Cytosine arabinosine 2mM (Sigma-Aldrich) was added at 8 days in vitro (DIV). For transfection using lipofectamine 2000 (Thermo Fisher), transfection was performed on DIV 8, using 1ml of lipofectamine and 1μg DNA in 1ml of medium per 35mm imaging dish, and medium was exchanged 6 hours later. Neurons were maintained in a humidified incubator with 5% CO_2_ at 37°C and imaged after 13-21 days in vitro. All experiments were performed on at least 2 independent cultures.

### cDNA constructs

To generate SSF-STANT in PCAGGS-SE, SSF-STANT sequence was amplified by PCR and inserted into pCAGGS-SE after digestion and ligation using EcoRI and XhoI sites. To generate SEP-DOR, SEP and DOR sequences were amplified by PCR and inserted into PCAGGS-SE using EcoRI and XhoI sites with a two fragments In-Fusion (Takara Bio) strategy. SSF-DOR S/T to A in PCAGGS-SE was generated with two fragments In-Fusion cloning in the NheI site. First fragment was a PCR amplification of the DOR sequence, and the second a codon optimized gene block that encodes the C-terminus tail of DOR with S and T amino acids sequences mutated to encode for A. SEP-DOR S/T to A was generated by PCR amplification of SEP and DOR S/T to A sequences and insertion into PCAGGS-SE using NheI sites with a two fragments In-Fusion strategy. All constructs were verified using sequencing, sequences for primers and gene block can be found in the extended methods section.

### Widefield imaging

Imaging of VAMP2-SEP for measurement of presynaptic activity, as well as imaging of SEP-DOR S/T to A for endocytosis was performed with a S Fluor 40x 1.30NA objective on a Nikon TE-2000 inverted microscope. System was controlled by Micromanager 1.4.10 software equipped with an Andor iXon EM+ EMCCD camera, and a Bioptechs objective warmer. Illumination, perfusion and electrical field stimulation devices were custom built (see extended methods and files on the repository https://doi.org/10.7272/Q63776Z3). Briefly, an insert was 3D printed and equipped with two platinum wires distant of 1cm used for electrical field stimulation (100 action potential at 10Hz, 10V/cm). Illumination in the green channel (VAMP2-SEP, SEP-DOR S/T to A) was controlled and synchronized with an Arduino uno, with a blue LED replacing the mercury bulb of a Nikon lamp and an appropriate combination of filters and dichroic mirror. Illumination in the red channel was achieved with a 543nm HeNe laser (Spectra Physics) and a micrometer-guided illuminator (Nikon), and syp-mCh fluorescence imaged with an appropriate set of filters and dichroic mirror. Neurons in glass bottom culture dishes were transferred from the incubator onto the system, culture medium removed and exchanged for HEPES buffered saline solution (HBS) imaging solution. HBS contained, in mM: NaCL 120, KCl 2, CaCl2 2, MgCl2 2, Glucose 5, HEPES 10 and osmolarity was adjusted to 270mOsm and pH to 7.4. This insert left a dead volume of 300μl inside the imaging dish and was used to perfuse solutions with a debit of 1.5ml/min.

To measure presynaptic inhibition neurons were nucleofected with VAMP2-SEP and opioid receptors. 2 minutes acquisitions at 1Hz were performed sequentially and electrical field stimulation starting at frame 59. First acquisition was always in HBS only to obtain a baseline response, ammonium chloride solution (HBS containing 50mM NH_4_Cl with NaCl adjusted to 80mM) was added with a pipette for the last 20 frames of the last acquisition. To monitor stability of the recording in SSF-MOR expressing neurons (“control”, figure 1C), the second acquisition was started 1 minute after the end of the first one and was in HBS only. To monitor acute inhibition at MOR, solution was shifted to HBS + DAMGO (Sigma-Aldrich) 10μM after the end of the first acquisition and the second acquisition was started 1 minute later. To monitor tolerance, neurons were subject to the same protocol except that they were pretreated with DAMGO 10μM directly in the cell culture medium for 18 hours in the incubator before imaging. When assessing desensitization after tolerance, the protocol was the same except that neurons were kept in DAMGO for 8 more minutes at the end of the second acquisition, and a third acquisition was performed still in the presence of DAMGO. For β-CNA experiments the protocol was the same as for measure of desensitization after induction of tolerance except that neurons were treated with β-CNA 50nM for 5 minutes and washed 3 times with HBS before imaging, not incubated with DAMGO. To obtain time course inhibition neurons were imaged as described when measuring tolerance except incubation time changed, and for the β-CNA conditions neurons were incubated with β-CNA 50nM for 5 minutes and washed 3 times with media before incubation with DAMGO 10μM for the time indicated. Acute inhibition and tolerance measurement for neurons nucleofected with VAMP2-SEP and SSF-MOR S/T to A and SSF-STANT was assessed as described for SSF-MOR. For measure of tolerance in the presence of Cmpd101 neurons were incubated with Cmpd101 30μM (HelloBio) or DMSO control in the culture medium in the incubator for 10 minutes before adding DAMGO 10μM for 18 hours and imaging as described previously. Tolerance to morphine 10μM (Sigma-Aldrich), PZM21 10μM and TRV130 10μM (generous gifts from Aashish Manglik, UCSF) with either Cmpd101 30μM or DMSO vehicle was assessed as described with DAMGO. Measure of acute inhibition, acute desensitization, tolerance, Cmpd101 effect for SSF-DOR and SSF-DOR S/T to A was assessed as described for SSF-MOR except the agonist used was DADLE 10μM (Sigma-Aldrich). To monitor the lack of cross tolerance between SSF-DOR and SSF-MOR, both receptors were nucleofected together with VAMP2-SEP and DAMGO 10μM or DPDPE 10μM (Sigma-Aldrich) added to the culture medium for 18h in the incubator before imaging. Neurons incubated with DAMGO were imaged in HBS, solution was changed to HBS + DPDPE 10μM and a second acquisition started one minute later, last acquisition was started 1 minute after switching the solution to HBS + DAMGO 10μM. Same was done for neurons incubated with DPDPE except the order of solution exchange was switched.

To monitor SEP-DOR S/T to A endocytosis, neurons were nucleofected with syp-mCh and SEP-DOR S/T to A and imaged in HBS for one frame in the syp-mCh channel and one frame in the SEP channel. Perfusion was switched to ammonium chloride solution and a second image taken 1 minute later. Perfusion was switched to HBS + DADLE 10μM and a third image taken 30 minutes later, the fourth frame was taken after 1 minute of perfusion with ammonium chloride solution.

### Oblique illumination imaging

Imaging of surface labeled opioid receptors at retromer marked endosomes, insertion events as well as quantification of surface fluorescence of SEP-tagged opioid receptors in axons was performed on a Nikon Ti-E TIRF microscope controlled by NIS-Elements 4.1 software. Microscope was equipped with an Andor iXon DU897 EMCCD camera, a perfect focus system and an objective and stage heater set to 37°C. Oblique illumination was achieved with 488, 561 and 647nm solid-state lasers (Keysight Technologies) coming at an oblique incident angle from an Apo TIRF 100x 1.49NA objective, and all channels imaged with an appropriate set of dichroic mirror and emission filters.

To image the recruitment of surface labeled receptors at endosomes, neurons that were transfected with VPS29-GFP, syp-mCh and SSF-tagged opioid receptors using the lipofectamine method were incubated for 15 minutes with Alexa-647 conjugated anti-FLAG antibody (1/1000, M1 antibody from Sigma, Alexa Fluor 647 Protein Labeling Kit from Thermo Fisher) before neurons were washed 3 times with HBS and mounted on the microscope in HBS. One image was acquired in the syp-mCh channel, rest of the acquisition was one frame in the Alexa-647 channel and one frame in the GFP channel every minute for a total length of the time lapse of 35 minutes. Agonist was added by pipetting 100μl of agonist-containing solution into the glass bottom dish after 5 minutes of baseline to a final concentration of 10μM, as indicated in figure legends. When neurons were treated with Cmpd101 or DMSO, neurons were incubated with Cmpd101 30μM or DMSO vehicle together with M1-Alexa647 1/1000 for 15 minutes. Cells were washed 3 times with HBS and mounted on the stage with HBS + Cmpd101 30μM or HBS + DMSO vehicle, rest of the protocol was the same as described previously.

For imaging of insertion events, striatal neurons were transfected using the lipofectamine method with SEP-tagged opioid receptors and syp-mCh. Cells were incubated for 20 min in the culture medium in the incubator with DAMGO 10μM (for SEP-MOR) or DADLE 10μM (for SEP-DOR), before 3 washes with HBS and mounting on the microscope stage in HBS + agonist at 10μM concentration. Cells in the no agonist conditions were washed in HBS and mounted in HBS without agonist. One frame was acquired in the red channel, and the green channel was imaged at 10Hz in stream mode for 5 min.

For imaging of surface fluorescence from SEP-tagged opioid receptors, neurons nucleofected with SEP-tagged opioid receptor and syp-mCh were washed 3 times with HBS and mounted onto the microscope stage in HBS. 30-40 random regions of interest were selected per dish in the syp-mCh channel only (experimenter was blinded to the green channel), and one image acquired in the green and red channel for each region. When neurons were chronically treated, cells were incubated for 18 hours with agonist at a concentration of 10μM in the culture medium in the incubator, the rest of the protocol was the same. If Cmpd101 or DMSO vehicle were present, Cmpd101 30μM or DMSO vehicle were added to the culture medium for 10 minutes before agonist 10μM was added for 18 hours, rest of the protocol was the same. Experiments were performed on the same culture in parallel for the conditions that are presented on the same graphs.

### Image analysis

Image analysis was performed on unprocessed 16 bits TIF images using custom written scripts in MATLAB (Mathworks, R2019b). Scripts are provided on the repository at the address: https://doi.org/10.7272/Q63776Z3. See extended methods for details of the procedures.

For the quantification of presynaptic activity, putative synapses were detected using an automated image classifier on an average baseline (defined as 6 frames before stimulation) subtracted ammonium chloride average image (defined as 5 last frames of the last acquisition). Fluorescence was quantified in a 5 pixels radius circle around each synapse, baseline fluorescence for each acquisition subtracted and normalized to the fluorescence in ammonium chloride. Synapses were validated if no pixel was saturated and the amplitude of the response during the first acquisition (in HBS only) was 5 times greater than the standard deviation of the baseline. For each condition and each acquisition, fluorescence from validated synapses were averaged across conditions and displayed in the inset curves (± standard error of the mean). To calculate the degree of inhibition, only acquisitions that had more than 50 validated synapses were considered and their fluorescence averaged across each acquisition. The ratio of the amplitude (defined as the maximum value of the 5 frames after stimulation) over the amplitude of the 1^st^ acquisition was used to define the degree of inhibition.

For the quantification of SEP tagged opioid receptors and syp-mCh fluorescence at synapses, synapses in focus were identified based on syp-mCh signal and manually picked, only exceptions were synapses that were over glial background autofluorescence or overlapping with somato-dendritic SEP signal. For each synapse, background subtracted average fluorescence within a 3 pixels radius was calculated for both channels. Values for each synapse and channel were normalized by the median fluorescence of the “no-agonist” condition of the same experimental day and data pooled between experiments.

To quantify receptor recruitment at retromer marked endosomes, a mask of VPS29-GFP endosomes was generated for each frame of the time lapse. To do so, regions of interest were manually selected on the image and refined by thresholding a maximal temporal projection of the receptor channel. Endosomes within this region were defined by a VPS29-GFP fluorescence value above a threshold set manually and objects larger than 5 pixels. For each region, background subtracted receptor fluorescence was calculated for each segmented endosomes for all frames. Average receptor fluorescence was calculated at all segmented endosomes of the same region, for the “no agonist”, baseline bin value. Receptor fluorescence at each endosome was normalized by this baseline value and averaged across all regions for 5 minutes intervals as indicated in the figures. Binned values were averaged across acquisitions for the same condition.

To quantify the frequency and fluorescence of single insertion events, bursts of fluorescence were manually selected on the image series. Fluorescence was quantified in a 2.2 pixels radius circle and the average baseline fluorescence of the 10 preceding frames was subtracted. Amplitude of fluorescence events is defined as the maximal fluorescence in the 10 frames following the detection. Fluorescence curves were averaged for all events of the same conditions. Frequency of events were normalized for each acquisition by the number of syp-mCh marked synapses in the field of view.

To quantify SEP-DOR S/T to A endocytosis using ammonium chloride unquenching, the 4 images were manually aligned to the first image to compensate for drift over the acquisition and lines drawn on axons identified based on syp-mCh signal. Background subtracted average fluorescence from a 3 pixel wide linescan was calculated for all images and normalized to the fluorescence value of the first image for each acquisition.

### Data presentation and Statistics

Quantifications of data are presented as either mean ± standard error of the mean, cumulative frequency curves, paired measurements or box and whisker plots (4 quartiles with inclusive median, outliers are not displayed, average is marked as a “X”). Graphs were generated with Excel (Microsoft office, 2016). Look up tables used can be found in the extended method section. All experiments were performed on at least 2 independent neuronal cultures and sample size indicated in the figure legends. When performing two tailed Student’s t-test the software used was Excel, two samples Kolmogorov-Smirnov test were performed with MATLAB. 95% confidence intervals were estimated by calculation of the mean over 50,000 random bootstraps using MATLAB.

