## Supplementary Information for "Endocytic trafficking determines cellular tolerance of presynaptic opioid signaling"

### Supplementary Information and Extended Methods

#### Legends for movies S1-S2-S3

Upper left panel is the syp-mCh image taken before the time lapse.

Upper right panel is the anti-FLAG M1-Alexa 647 image series (receptor).

Bottom right panel is VPS29-GFP image series (endosome).

Bottom left panel is the segmentation of endosome signal on which the quantification is performed. Dark red is manually selected polygon, light red is the refined region of interest (thresholding of the maximal projection of the receptor channel), white is segmented endosomes (thresholding of VPS29-GFP signal within the refined mask).

Full frames are shown for each panel (~80μm*80μm), total length of the time lapse is 35min imaged at 1 frame per minute, movie is 10 frames per second.

Movie S1: SSF-MOR internalization in axons. Same as for figure 2A, DAMGO 10μM is added after frame 6.

Movie S2: SSF-DOR internalization in axons. Same as for figure 4A, DADLE 10μM is added after frame 6.

Movie S3: SSF-DOR S/T to A internalization in axons. Same as for figure 6D, DADLE 10μM is added after frame 6.

#### cDNA constructs:

Primers for SSF-STANT in PCAGGS-SE:
SSF-STANT forward: GCGCCTCGAGATGAAGACGATCATCGCCCTGAGC
SSF-STANT reverse: GCGCGAATTCTTAGGGCAATGGAGCAGTTTCTGC

Primers for SEP-DOR in PCAGGS-SE using In-Fusion two fragments:
SEP forward: ATATCGGTACCTCGAGATGGACAGCAAAGGTTCG
SEP reverse: CCAGCTCCATGGTTTTGTATAGTTCATCCATGCC
DOR forward: AAACCATGGAGCTGGTGCCCTCTGCCCG
DOR reverse: CCTGAGGAGTGAATTCCTCTAGATTATCAGGCGGCAGCGCCACCGCCC

Primers and gene block for SSF-DOR S/T to A in PCAGGS-SE using In-Fusion two fragments:
SSF-DOR forward: ATATCGGTACCTCGAGCTAGATGAAGACGATCATCGCCCTG
SSF-DOR reverse: CGACAGAGCTGGCGGAAGCAGCGCTTGAAGTTCTCGTCCA
Gene block:
TGCTTCCGCCAGCTCTGTCGCGCCCCATGTGGGCGACAAGAGCCTGGTGCTCTCAGAC
GACCCAGACAAGCAGCTGCTCGAGAACGCGTAGCAGCATGCGCACCAGCCGACGGGC
CAGGGGGTGGGGCTGCCGCATAACTAGCGGCCGCATGCGAATT

Primers for SEP-DOR S/T to A:

SEP forward: CGATATCGGTACCTCGAGCTAGATGGACAGCAAAG
SEP reverse: GGCACCAGCTCCATGGTTTTGTA
DOR S/T to A forward: ACAAAACCATGGAGCTGGTGCCC
DOR S/T to A reverse: AATTCGCATGCGGCCGCTAGTTATGCGGCAGCCCCACCCCC

#### Images look up tables:

low Intensity high intensity

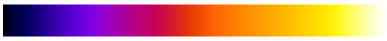
SEP-MOR, SEP-DOR (Fig 6F), SEP-DOR S/T to A (Fig 6F)

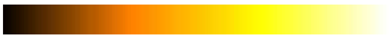
VAMP2-SEP (Fig 1B), syp-mCh (Fig 2A, 4A, 5B, 6F)

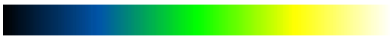
VPS29-GFP

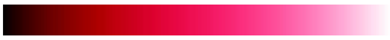
M1-alexa647 labelled SSF tagged opioid receptor

#### Custom built hardware and acquisition protocol:

##### Insert for perfusion and stimulation

A custom made insert for matTek 35mm glass bottom dishes with 14mm coverslips (P35G-1.5-14-C) was designed and 3D printed on a uPrint plus (Stratasys) from AMS plastic, the .stl file necessary for this part is provided. We recommend the use of black material for the insert to minimize artifacts of fluorescence (Fig 1A). A rubber O-ring (15mm internal diameter, 1mm thickness) is installed at the bottom of the insert, these are standard and can be found at any hardware store (Fig 1B). After 3D printing of the part and extensive washing with water (typically a day in >500ml water with multiple replacement of the water, to wash out any residual plastic and alkaline solution), stimulation wires were setup on the insert. We recovered platinum wires (2x1.5cm) from a broken electrophoresis device (Fig 1C) and soldered to about 5cm regular electrical wires (2 per insert) using a soldering iron (Fig 1 D,E). Starting from the platinum end, the wire is inserted into a cut 10ul pipet tip. The platinum wires will stick out of the tip opening while the solder and regular wire would stay inside, we cut another 10ul pipet tip and inserted it in the one containing the electrical wire to squeeze it and secure the wire in place (Fig 1F,G). The platinum wire is bent 90 degrees from the cut pipet tip and inserted into the appropriate semi-open cylinder, repeat on other electrode (Fig 1 H,I). Using small tweezers, shape the platinum wire so it follows the bottom of the insert and secure the end into the appropriate hole in the insert (Figure 1 K,L). The stimulation wires are distant of 1cm once set up. We printed a dish holder that fits the stage of the microscope, it holds 35mm dishes and accommodates a rubber band (run it over the middle of the opening), the .stl file for printing is provided (Fig 1L).

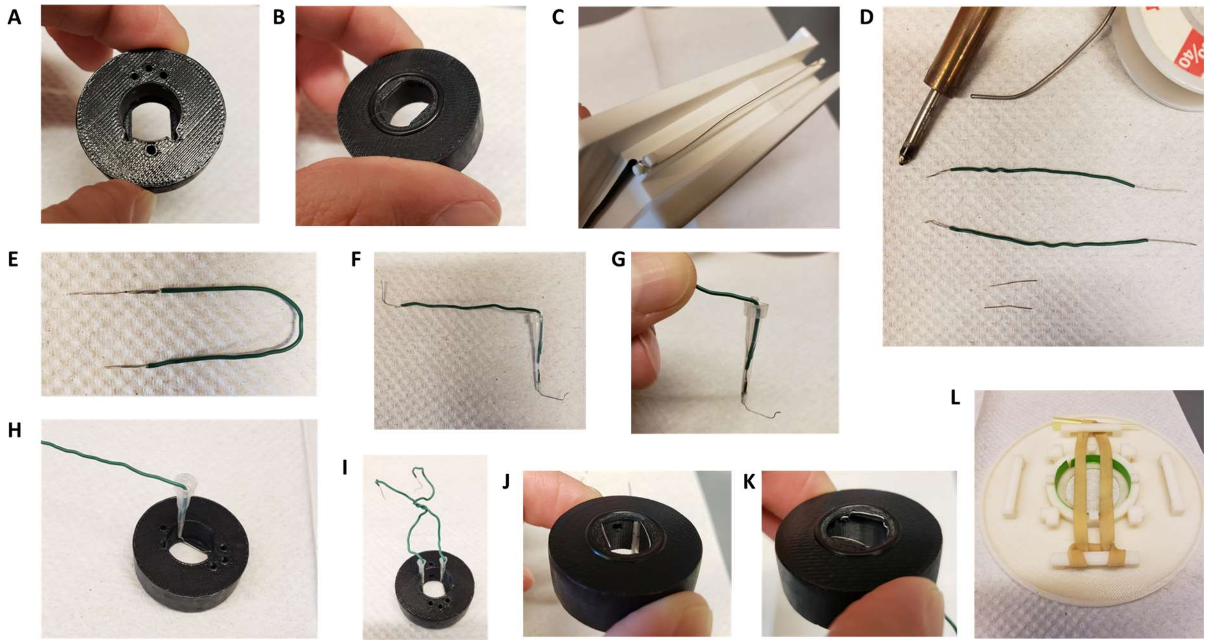


**Figure1:** Insert for perfusion coupled to electrical stimulation. **A.** Top view of the insert, with 3 holes for perfusion entry in the insert (top), one hole for vacuum suction (bottom), and 2 semi-open holes for electrodes (bottom). **B.** Bottom view of the insert with O-ring in place. **C.** Broken gel electrophoresis with platinum wire. **D.** Electrodes are made by soldering 2 platinum wires (bottom) to regular electric wire. **E.** Platinum wire soldered to the electric wire. **F.** Electrode secured and ready to be installed. **G.** Close up view of the electrode, note the angle. **H.** Insert with one electrode installed. **I.** Insert with both electrodes installed, we recommend you twist the electrical wires together to solidify the installation and make sure the hole for vacuum stays easily accessible. **J,K.** Bottom view of the insert with the two electrodes installed. **L.** Dish holder for 35mm dishes with rubber band installed.

<u>Zapomatic</u>

**WARNING**

**DO NOT ATTEMPT ANY WIRING OF ANY KIND if you lack the knowledge and understanding required. Electricity is dangerous and can cause personal injury or DEATH as well as other property loss or damage if not used or constructed properly. If you have any doubts what so ever about performing do-it-yourself electrical work, PLEASE do the smart thing and hire a QUALIFIED SPECIALIST to perform the work for you. Always disconnect the power source before working with electrical circuits. This information is provided for the use of individuals as they see fit! All parties associated with it are not responsible for the use and results of this information by any party, especially those lacking sufficient skill or knowledge to perform these steps safely and ANY HAZARD CREATED IS THE SOLE RESPONSIBILITY OF THE USER.**

This device can is built from basic hardware and an Arduino UNO circuit board (see details of the electrical circuit provided in figure 2). The zapomatic needs 30V electrical supply. The electrical wires of the insert are connected to the zapomatic using a BNC cable equipped with a pair of alligator test clips (see Fig 4G).
The Arduino code provided “Dam100AP” needs to be uploaded on the Arduino. The board receives a TTL high signal from the camera or illumination and counts the number of frames. Once a certain number of frame is reached (defined by user, here 59), the zapomatic will fire electrical pulses through the BNC (length of each pulse/frequency/number of pulses user defined, here 100x 1ms pulses at 10Hz). Intensity of electrical stimulation is adjustable manually on the device, and was calibrated to 10-12V/cm using an oscilloscope connected in parallel from the stimulation BNC output (distance between the electrodes is 1cm). The zapomatic counter is reset after the firing sequence, and zapomatic will fire again any time it reaches the count threshold (just once with our acquisition setting). A manual button on the device can be pushed to trigger firing manually. A LED lights up when the device is firing (same frequency/duration).

If LED light is dim when firing the current does not flow between electrodes. Oxidation can build up at the solder point, check for proper conductivity.
Frame count will reset to 0 after 10s without TTL high.
If counting frames directly from the camera TTL output we recommend you disconnect the input to zapomatic or turn off the firing switch until ready to start the acquisition.

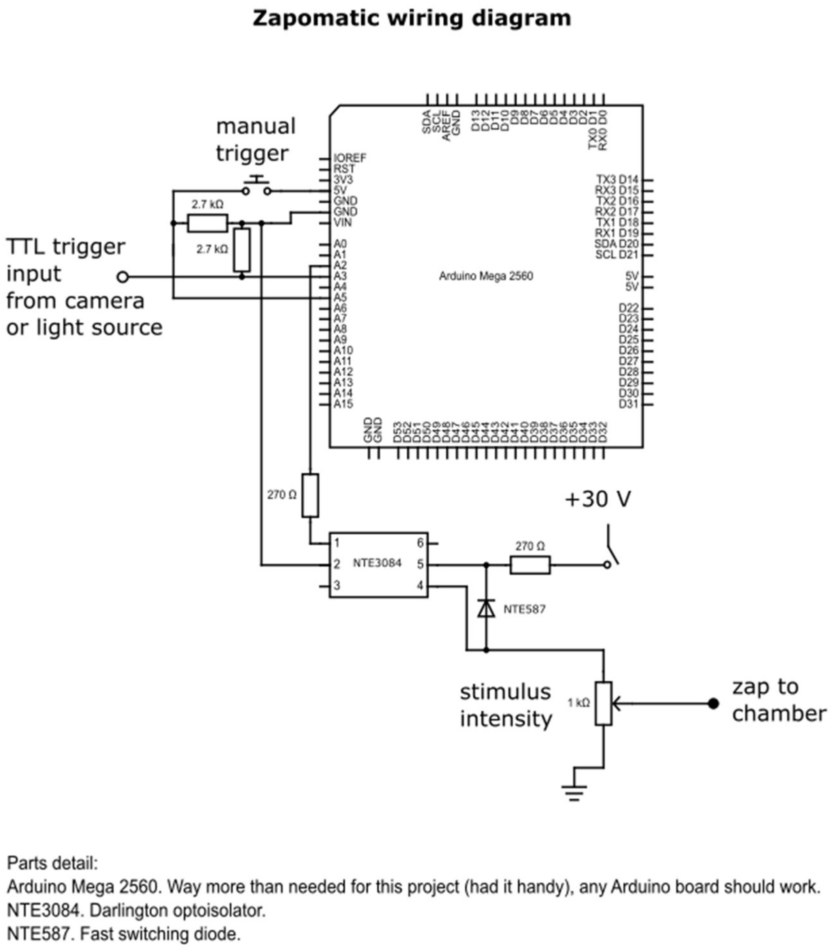


**Figure 2:** circuit diagram of the zapomatic

Setting up the insert:

The perfusion is based on gravity flow from up to 3 different solution sources, each line consisting of an assembly of (from top to bottom): syringe (containing the solution), a 0.45µm filter, a manual valve, tubing, and ending with a 10ul pipet tip (Fig 3). A separate vacuum line to remove solution need to be setup and attached to tubing ending with a 10ul tip. We used two 30ml syringes filled up with ~35ml of solution. One contained control imaging solution (HEPES buffered saline solution (HBS) adjusted to pH 7.4 containing, in mM: NaCL 120, KCl 2, $CaCl_2$ 2, $MgCl_2$ 2, Glucose 5, HEPES 10 and osmolarity was adjusted to 270 mOsm), the other HBS + drug (typically DAMGO or DADLE at 10µM). Syringes are placed vertically so the filter is at about the height of the stage (Fig 4H). When setting up, use the syringe piston to suction the air trapped in the filter (valve closed), load the whole line with solution, and remove any air bubble. When opening the manual valve solution flows from the end of the line by gravity, adjust the differential in heights between the top of the solution in the syringe and the insert to adjust the flow. In this setting the perfusion runs at about 1.5ml per minute and we readjust the volume between acquisitions. See figure 2 for a schematic of the whole setup.

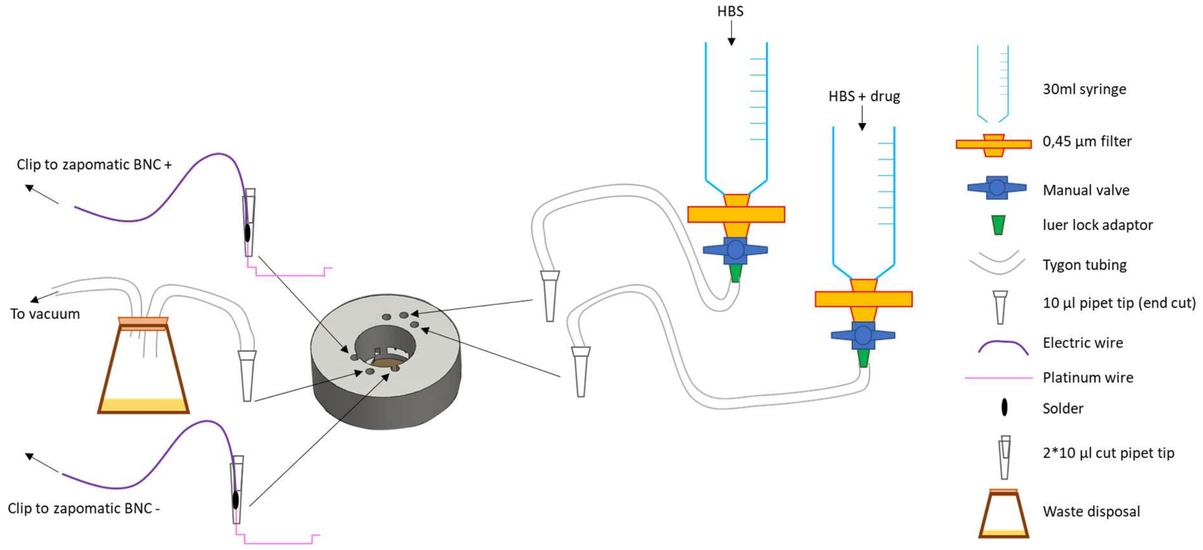


**Figure3:** schematic of the perfusion installation.

If the vacuum does not work efficiently there is a risk of overflow and damage to the microscope.
The mounting of the dish is done on the microscope stage but away from the optics.

Primary neurons were cultured in matTek 35mm glass bottom dishes with 14mm coverslips (P35G-1.5-14-C, Fig 4A). Insert is mounted into Using Molykote vacuum grease (Motion industries, # 00785543) inside a syringe (Fig 4B), create a uniform ~2mm thick layer of grease surrounding the O-ring (Fig 4C). The dish is brought to the microscope from incubator (37°C 5% CO2) and media inside the dish removed as extensively as possible while not approaching the suction close to the glass part to prevent drying of the live cells (Fig 4D). The plastic part of the dish is further dried using a rolled kimwipe (important to make a good seal), and the insert pressed onto the dish. Do not release the pressure until the end of installation. Setup the perfusion lines in the appropriate entries and open the valve for the vehicle solution (Fig 4E). Wait for it to flow above the platinum wires and place the vacuum line in the appropriate hole. The insert is designed for strong suction such as a vacuum pump connected to a waste container. Suction should be steady and volume inside the insert constant, weak suction can result in changes in volume of solution inside the insert which cause focus issues. This insert left a dead volume of 300ml inside the imaging dish. Once the perfusion and vacuum lines are setup, place the rubber to secure the dish and place the dish holder on the stage (Fig 4F), and use the alligator test clips to connect the electrodes to the zapomatic (Fig 4G). If using a different dish holder, we recommend you clamp the insert on the dish to prevent movement. The acquisition is ready to start, make sure the solution lines are never empty. At the end of the acquisition, remove the solution and vacuum lines from the insert, place the ruber bands on the blocks away from the insert, and pop the dish out of the holder by pressing the dish holder on something into the objective opening. Gently take the insert out of the dish being extra careful to not displace the electrodes and use scalpel or any flat blade to remove the grease from the insert (Fig 4I), finish cleaning the bottom of the insert with a kimwipe. After each day of acquisitions, the perfusion lines are abundantly rinsed with distilled water (typically >150ml per line) with the insert and vacuum line in place in the last imaged dish. Using the plunger all lines are emptied from remaining water (pump air into the filter/tubing, remove syringe from the line, load air into the syringe, couple to line, push the plunger. Repeat a few times).

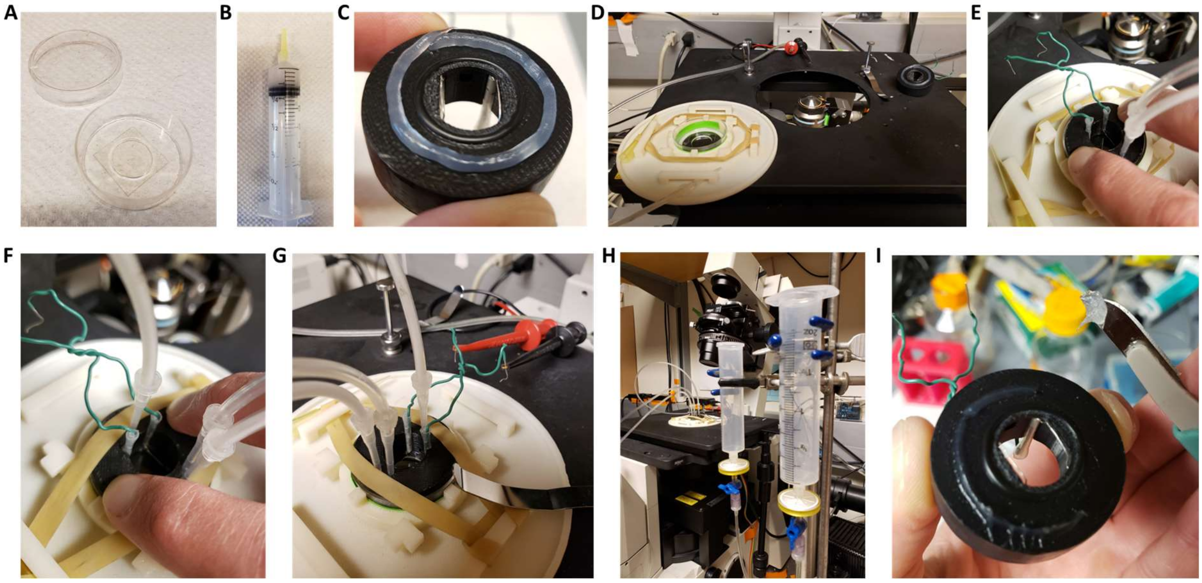


**Figure 4:** Setting up of the insert. **A.** 35mm glass bottom dish with 14mm coverslip, note the plastic area surrounding the coverslip that needs to be completely dry for the vacuum grease to create a good seal. **B.** Vacuum grease loaded into a 30ml syringe completed with a cut pipet tip. **C.** Grease applied to the bottom of the insert. **D.** Dish setup in the holder with the insert ready to be installed. **E.** Insert with perfusion lines installed, keep pressure on the insert to maintain a good seal. **F.** Vacuum line installed and rubber band securing the insert, after this step it is no longer necessary to keep pressure on the insert. **G.** Dish older with insert installed on the optics and electrodes connected to the alligator test clips. **H.** View of the whole system in place. **I.** After removing the insert use a scalped to remove the excess grease for the bottom of the insert, finish cleaning with a kimwipe.

<u>Microscope and acquisition software:</u>
Imaging of presynaptic activity in striatal neurons that were nucleofected opioid receptors together with VAMP2-SEP was performed on a Nikon TE-2000 inverted microscope, objective was a S Fluor 40x 1.30 NA objective, an Andor iXon EM+ EMCCD camera and a Bioptechs objective warmer. Widefield illumination in the SEP channel was controlled and

synchronized with an Arduino uno, with a blue LED replacing the mercury bulb of a Nikon lamp and an appropriate combination of filters and dichroic mirror. Acquisition software used was micro manager Version 1.4.10.

Acquisition settings:

For most experiments described in this study, the acquisition protocol was standard and consisted in two acquisitions of 120 frames a 1Hz (second acquisition usually started 1min after switching the valves from vehicle solution to vehicle + drug). Camera was encoding images at 14 bits and saved as 16 bits 512*512 .tif images. The Arduino code embedded in the “zapomatic” triggered 10Hz – 1ms stimulation starting at frame 59 for a total of 100 stimulations – 10s. Control imaging solution was then switched to imaging solution + drug by switching valves on syringe assemblies (open first and close second to ensure continuous flow of solution) for one minute before starting the second acquisition. At the end of the second acquisition, 1ml of a solution containing ammonium chloride (HBS containing 50mM NH4Cl with NaCl adjusted to 80mM) was pipetted in the open chamber after closure of the perfusion valve, starting at frame 100.

Time in presence of agonist or sequence of HBS + drugs varied across experimental designs as described in the manuscript. When a third acquisition was performed NH4Cl solution was pipetted at the end of the third acquisition.

#### Analysis of presynaptic activity:

All the workflow presented here is setup for analysis in batch (software handles multiple acquisitions at the same time). User will need to install the different scripts provided and add them to the path in MATLAB. In addition, it requires the image processing toolbox and the parallel processing toolbox (optional but slower, see notes). Sample data are available for running the analysis.

Detection of synapses:

Manual mode:

Set the current folder containing the last movie of the acquisition with NH4Cl at the end (“Acquisition#” in the sample dataset). In the command window, enter “**zap100_1_2**”. Program will ask to select the corresponding file (.tif file ending in -2_1.tif in the sample dataset) and a graphical user interface will appear:

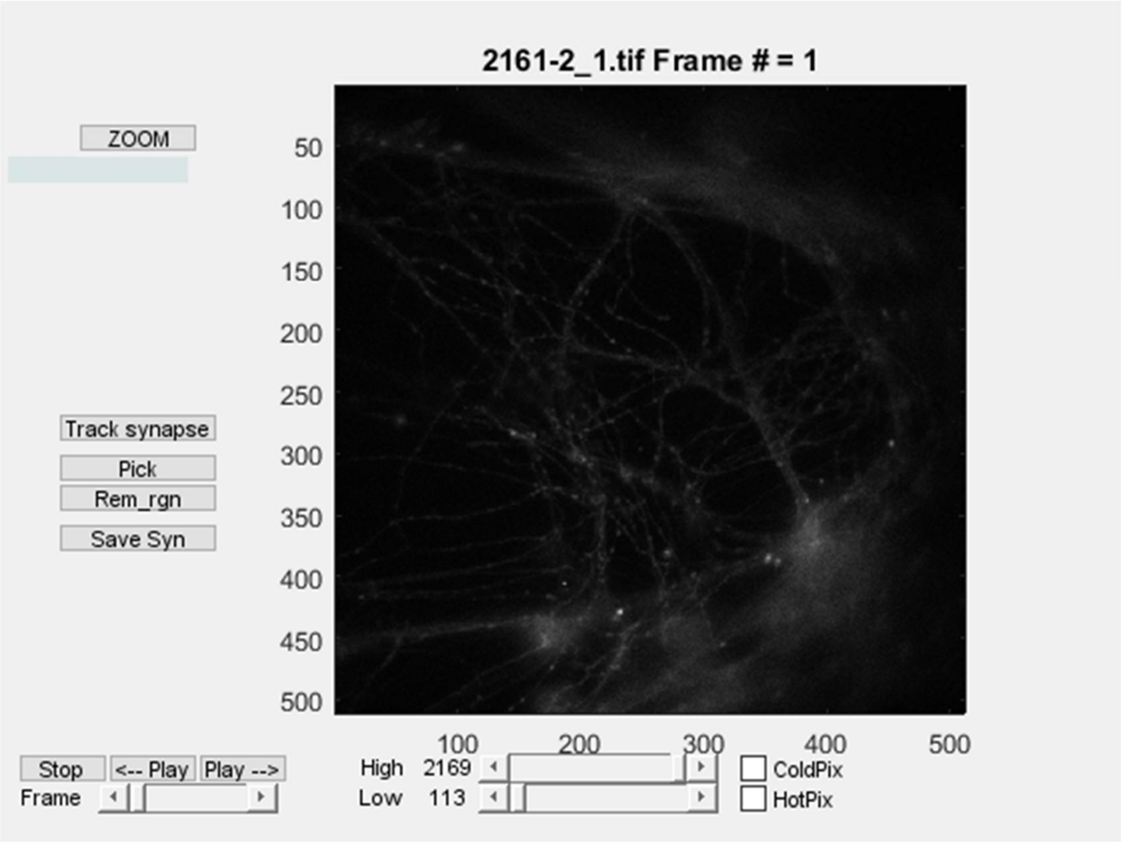


**Figure 5:** Zap100_1_2 user interface. Title is name of the file, # indicates the frame (here 1). Lower list of buttons (Track synapses, Pick, Rem_rgn, Save Syn) are controls to select synapses (see below). Lower left buttons are controls for the frame and play the movie. Middle lower controls are to set the contrast.

The manual workflow is as follow, Review the movie for quality first. When clicking on "track synapses", select "yes" to manually pick, and program will ask to identify a background region (rectangular), and will automatically go to frame 121, the last one. This frame does not belong to the original movie but instead is a differential image of the last 5 frames of the movie (in NH4) minus an average of the 10 frames that precede the stimulation + 8000 (AU). These parameters can be found 27-31 of the code. Synapses are much easier to identify on this image. Any left click on the image will select a synapse, that appears as a green circle. Right click will stop this sequence. You can select more synapse anytime (after zooming if needed) or remove synapse one at a time using the "pick" button. Right click when "pick" button is activated removes synapses. Use Rem_rgn to remove multiple synapses within a rectangular region. When satisfied with the selection, save the list of synapses using "Save Syn". It will generate a .txt file that ends with ZAP1_2_syn.txt that contains the coordinates of synapses as follow:

| | |
|---|---|
| 59 5 10 5 | parameters for the diff NH4 image |
| 69 325 46 15 | coordinates of background region |
| 1 120.78 160.6 0 | synapse 1 X Y coordinates |
| 2 163.04 171.98 0 | synapse 2 X Y coordinates |
| 3 200.42 165.48 0 | synapse 3 X Y coordinates |

And so on…

Using machine learning:

**Step1: generate sample synapse images.**

For generating the sample images it is important that you select ALL synapse-looking structure on the image when selecting them manually. Anything on the image that isn't registered as a synapse will be treated as background by this code. Organize your current folder as follow (as organized in the sample dataset, standard for all batch processing), that usually reflect multiple acquisitions (folder 1.1,1.2 etc.) for multiple conditions (folder 1, folder2):
Code as setup will recognize the movie with NH4Cl at the end because the name finishes by 2_1.tif (line 46). This is the way micromanager saves the second acquisition in our system. Either adapt code to a unique identifier for NH4 movie or add "2_1.tif " to the file name.

Folder 1 CONDITION FOLDER 1
  Folder1.1 ACQUISITION FOLDER 1.1
    .tif movie of first acquisition (acquisition 1)
    .tif movie of second acquisition with NH4 at the end (acquisition 1)
    .txt file with annotated synapses (acquisition 1)
  Folder1.2 ACQUISITION FOLDER 2.2
    .tif movie of first acquisition (acquisition 2)
    .tif movie of second acquisition with NH4 at the end (acquisition 2)
    .txt file with annotated synapses (acquisition 2)

  And so on…

  Folder1.3
    …

Folder 2 CONDITION FOLDER 2

  Folder2.1 ACQUISITION FOLDER 2.1
    .tif movie of first acquisition (acquisition X)
    .tif movie of second acquisition with NH4 at the end (acquisition X)
    .txt file with annotated synapses (acquisition X)
  Folder2.2 ACQUISITION FOLDER 2.2
    .tif movie of first acquisition (acquisition Y)
    .tif movie of second acquisition with NH4 at the end (acquisition Y)
    .txt file with annotated synapses (acquisition Y)

And so on
Folder 3
  …

Enter in the command window "**MLsynGen**"
A few parameters are embedded in the code, same ones used for the differential NH4 image (line 12-14) and "boxsize" (defines the size of each generated image, default is 7) and "samplesize" (how many images to generate from each acquisition, default is 50). As it is set up, code will randomly pick 50 synapses per acquisition at most (line 9) and will generate 3 stacks (it can take time depending on the amount of images you have, be patient):

Stack 1: sampleSyn.tif, the multi-tif stack contains synapse images so central pixel is centered around the manually selected coordinates, and a boxsize selection of the default NH4 image around it ( so by default, size of the stack is 15*15*n images, n depending on

samplesize or min amount of selected synapses, 15 being 2*boxsize +1).
Stack2: sampleEmpty.tif, , the multi-tif stack contains images selected randomly around synapses (this is a “close by” sample picked from a distance of boxsize away from manually picked synapses), size of the stack is 15*15*2n images.
Stack3: sampleBack.tif, the multi-tif stack contains images selected randomly in the background, size of the stack is 15*15*2n images.

Provided sample multi-tif stacks used to train the classifier used in this study have n = 20643 images.

**Step 2: train and run the classifier**

In the condition folder, enter “**HogClassBatchGaussAI**” in the command window.

Code will ask to select a synapse movie, pick sampleSyn.tif

Then to select “Off movies” 1 and 2, select sampleEmpty.tif and sampleBack.tif (order does not matter).

Code will train a support vector machine classifier that is fed a histogram of gradient vector after transform of the images (enlarge + gaussian filter + Fourier processing) and an intensity value (center – background around). All parameters are found line 21-26. It is trained for 2 classes based on the images provided, synapse (1) and not synapse (0). A confusion matrix is displayed once the classifier is trained, and classifier is automatically saved in the current directory, “HogFFTclass.mat”.

Classifier will analyze all provided acquisitions, this mode does not run on parallel processing and is slow. To detect synapses, a sliding window operates with a 1.5*boxsize (parameter divwind, 1.5 is default) step and generates an image that is filtered for signal intensity (uniformly high signal or nearby “negative value” that indicate movement are excluded, line 273) and classified as synapse or not synapse. If classifier found a synapse, it will operate a 2D gaussian fit to center the localization of the synapse. Parameters of the gaussian fit are found line 287-289 and will need to be adjusted depending on your acquisition setting (typically create an average projection of your “synapse” sample and extract parameter from this). Script will then remove double detections (<tooclose, 3 pixels is default).

For each movie it will generate a “MLsynHFA.txt” synapse coordinate file in the Folder X.Y of each acquisition. You can visually check the quality of the classifier by running Zap100_1_2 in the command window (select “no” to manually annotate, “yes” to load the background region and select original manual file, or select “no” and manually pick a background region). You can manually edit the synapses at this stage, and save the file as explained before. If classifier performance looks good you can use it directly, no need to re-train every time. The classifier used in this study is provided in the “sample for classifier” dataset.

All parameters provided with this code have been set through iterations and will vary depending on your imaging system (magnification etc.). Make sure your classifier gives great performance on selected data before generalization of its performance to new datasets (>95% accuracy on both classes as defined from classifier output).

**Step 3: run classifier on new data**

Organize the dataset to analyze in the current directory as described previously for CONDITION (multiple folders for each acquisition). Copy the classifier generated from step 2 into the current folder, enter “**ClassGaussAllign**” (**ClassGaussAllign3** if 3 acquisitions) in the command window. This code runs on parallel processing, but you can change "parfor" line 31 for a "for" statement and code will run. Classifier will detect synapses as explained previously and generate a "_CGA.txt" file that contains the synapse coordinates. User will need to select background region using ZAP100_1_2 (select "no" to manually annotate, "no" to load background region, and manually pick a background region). While I could have automated this step, I find it useful (and fast) to select the background region while checking quality of synapse detection and eventually adding – removing synapses manually.

<u>Quantification of fluorescence and normalization of the data:</u>

Set the current folder as described previously (and as orgamized in the sample dataset), with the multiple conditions folders and acquisitions subfolders (2 movies + 1 "ZAP" synapse + background coordinate file). Code looks for "ZAP" in the filename to identify the coordinate file, for "2_1.tif" to identify the second file with NH4 at the end, and pick the other ".tif" file as default for the first control movie. You will to rename the files or change the code accordingly (lines 92-94). Enter "**zap100Poolstdev**" in the command window ("**zap100Pool3stdev**" if 3 movies). Code requires "xlswrite" function that (to my knowledge) does not work on MAC OS version of MATLAB. User can convert the corresponding cells to tab format and write excel or csv file. I will expand here considering the 2 acquisitions and a note can be found for 3 acquisitions paradigm at the end. While a number of parameters are embedded in the code for the sake a more friendly user interface, I will provide line numbers for these.

Code will analyze each acquisition, normalize and pool the data for the whole condition, and normalize again. Details of the operations are found below:

<u>For each acquisition</u>, code will generate a .xlsx file containing multiple tabs
First tab contains the parameters, the defaults are:

- First stimulation: default 59, in our setting, marks the beginning of the stimulation
- Size of the synapse: default 5, radius in pixels around synapse coordinates in which the fluorescence measurement is done
- baseline: default 6, number of frames before stimulation to define fluorescence baseline
- Amplitude: default 5, number of frames to define the amplitude of fluorescence increase after stimulation
-Stimulation: default is 10, length of stimulation in frames.
-Frames in NH4Cl: default 5, to define fluorescence in NH4Cl.
-Threshold Amplitude: default 5, to select only responsive synapses.

How these numbers are used for quantification is explained below.

For each coordinates, the script will calculate raw fluorescence values (average fluorescence in a circle of radius "Size of synapse") and subtract the average fluorescence in the user defined background region. This appears in two tabs with "_raw" added at the end of the movie file name.

The script then normalizes the fluorescence values for each synapse. To do so it defines the baseline fluorescence as the median fluorescence value between the frames "first stimulation – baseline" and "first stimulation". Baseline fluorescence value is subtracted from raw values. Script will calculate the average raw fluorescence of the last 5 frames of the NH4 application, and this value is used to divide the baseline subtracted raw fluorescence values, for the two movies. This appears in two tabs with "_NH4norm" at the end, normalized fluorescence over fluorescence in NH4.

In these tabs there are two extra columns at the very end that are labelled "amplitude" and "STAMP". These values are only used for classifying synapses as responsive. Amplitude is the average of _NH4norm fluorescence between the frames (First stim + Stim) and (First stim + Stim + Amplitude), with the 20% highest and 20% lowest values excluded (line 247) to minimize noise. "STAMP" corresponds to this amplitude value divided by the standard deviation of the baseline _NH4norm fluorescence between the frames "first stimulation – baseline" and "first stimulation".

Based on these calculations, a number of synapses are excluded.
-Synapses that have values above the NH4 average value before NH4 is applied (20 frames, line 215).
-Synapses that contain 1 or more saturated pixel (value > 16300 based on 14 bits, line 215) during the last NH4 frames.
These two categories of synapses will appear in the tab "_removed" of the excel file with a value 1 for "saturation".

-Synapses for which STAMP < Threshold for the first acquisition, this is to remove structures detected as synapses that do not respond to the stimulation (low signal to noise). This will appear in the tab "_removed" of the excel file with a value 1 for "amplitude".

The two remaining tabs are labeled "_final" for each movie and contain the quantifications only for the synapse that passed these exclusion criteria.

For each condition, the script will pool data from this single-acquisition analysis and perform another normalization for the whole condition folder, the results are found in a "_std_Pool.xlsx" file.

All quantifications for synapses that have been validated are pooled into two tabs, one for each movie ($1^{st}$ Stim, $2^{Nd}$ Stim), with the name of the corresponding acquisition file in the first column. Last column represents the amplitude of the response as defined previously. Average curves +/- SEM presented in the manuscript were obtained from this dataset.

For each acquisition/movie, an average of fluorescence and amplitude for validated synapses is calculated, this appears in the tabs "$1^{st}$ cell, $2^{nd}$ cell", as well as the number of validated synapses for each acquisition "n". Reminder, the average amplitude here is not what was used for final quantification, we choose to quantify the amplitude of the average fluorescence - not the average of the amplitudes, to improve signal to noise. This is explained below.

For each acquisition, we therefore obtain two normalized over NH4 fluorescence curves for validated synapses ($1^{st}$ movie baseline, $2^{nd}$ movie + drug, usually). We here define the amplitude of the average as the maximum fluorescence value between the frames (First stim + Stim) and (First stim + Stim + Amplitude), that is calculated for each movie. This appears into the "final" tab of the excel file under "Max Amp", for each movie ($1^{st}$ STIM, $2^{Nd}$ STIM). Average fluorescence curves for each movie are normalized to the Max Amp value of the first (control) acquisition and displayed into this "final" tab, as well as averages and SEM value for the whole condition. To define the degree of inhibition in the present manuscript, we only considered acquisitions with at least 50 validated synapses and used the ratio such that the percentage of inhibition is given by:

100*(1-(Max Amp(2$^{nd}$ movie) / Max Amp (1$^{st}$ movie, control))).

In the 3 movies/acquisition paradigm, we used "**zap100Pool3stdev**" , with folder containing 1 baseline movie "1_1.tif", 1 drug movie "1_2.tif", 1 drug movie + NH4 at the end "1_3.tif", 1 synapse .txt coordinate file "ZAP". Script will process as previously for 2 acquisitions except that it quantifies another movie but all synapse validation and normalizations steps remain the same.

#### Analysis of receptor recruitment at endosomes:

The code runs by entering "**Manon**" in the command window of MATLAB.

If you run the code multiple times on the same movies, make sure that the previous files generated by the program are in a different directory or code will overwrite and edit.

**Code asks for "size of the structure",** this is the minimal size in pixels of the structure you would be segmenting. Default is 5, and this is what was used in the study.
**Code asks for synapse movie**, this should be a multi-tif single channel. Code does no operation on this image; it is for display only. If only two channels have been aquired, provide a movie of the same size (repeat marker or receptor for example). It will appear in blue in the image display.
**Code asks for marker movie**, this should be the movie you try to segment, it will appear in green in the image display.
**Code asks for receptor movie**, this is the signal you try to quantify, it will appear in red in the image display.

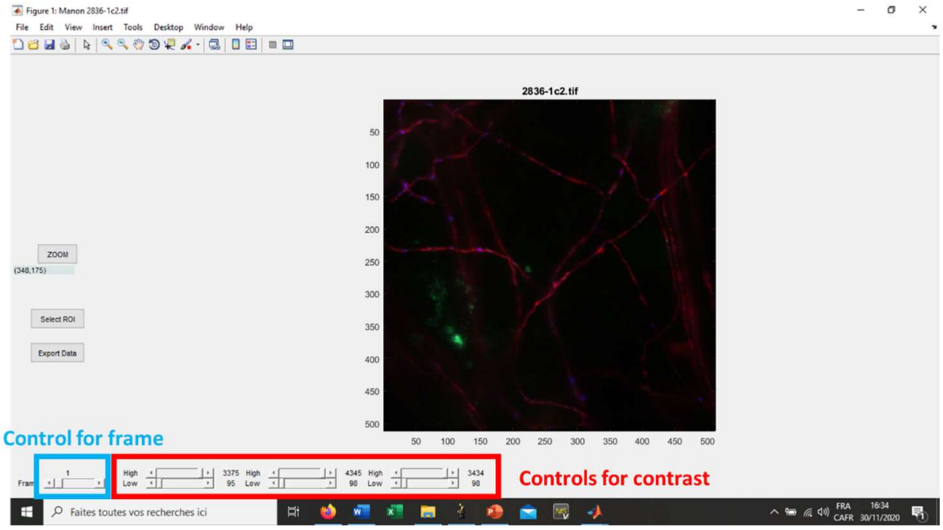


**Figure 6:** User interface for Manon.

This is the user interface for this analysis that will pop up.
If not: make sure all your movies are the same size (frame number in particular), make sure you set the path correctly (for the code and for your working directory), make sure you have the proper toolbox installed (image processing toolbox)

Set contrast and look at your images to check for quality using the controls. Zoom button can be used to zoom on some area of the image, use right click/reset to original view to get back to original image.

Quantification starts by clicking on “select ROI”.
Program asks to select a rectangular background region, draw the rectangle with the cursor and double click on the rectangle to validate the background region.
Program returns control of a cursor, use it to draw a polygon around a ROI (left click), double click to close and validate the polygon. Another interface will pop up:

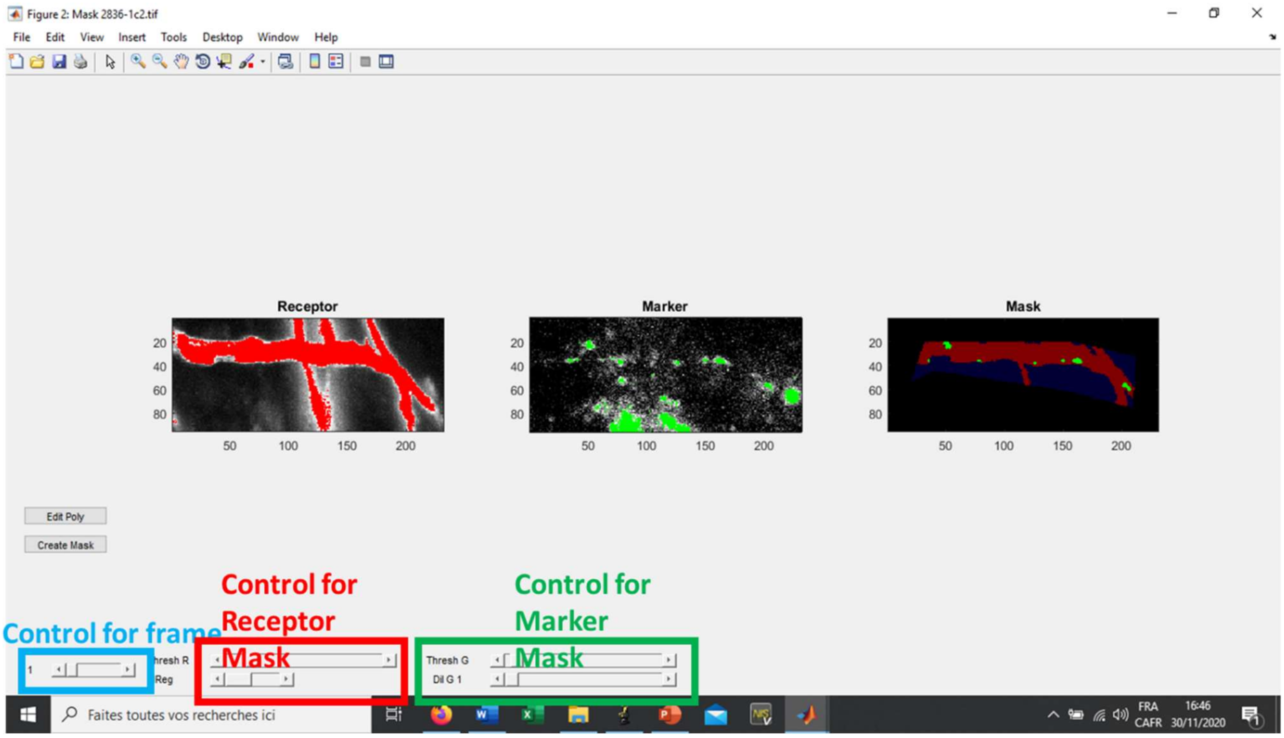


**Figure 6:** segmentation interface Manon.

Use the controls to define the segmentation:

The receptor mask is made on the maximal projection of the receptor movie (red, image on the left). User can control a threshold above which you define the mask, and the number of regions to include in the mask starting from the biggest. This mask appears in the “mask” window as red over blue background (the polygon ROI). This mask is used to refine the segmentation in the marker channel.

The marker mask is generated using two controls, I recommend you use the threshG control only that sets the threshold for the segmentation. Other parameter for segmentation would be the size of the object (inputed at the beginning of the code, 5 in this study) and dilG that lets you soothe the structures (1 in this study). The segmented structures within the receptor mask (middle, green) appears on the “mask” window as green dots over red background.

When satisfied with the segmentation, click on “create Mask”. The window closes and the polygon you drew appears stably on the image. If you’re not satisfied with the ROI you selected, click on “Edit Poly”, it will erase the polygon like nothing ever happened.

When you have selected all the ROIs on the image, click on export data. Leave some time for MATLAB to do its job as writing on excel takes a bit of time, script will display the message “done”, only then you can explore your results in the excel file. The file is added to the current directory and is called “name of your synapse movie_Manon.xlsx”. This file contains the quantifications:

In the first tab, every column is a frame of the movie.

**back Fluo** is the average fluo in the background region.
**Average Fluo Receptor** is the background subtracted average fluorescence of the receptor in the max projection mask, per ROI (multiple lines, per ROI).

**Average per region Fluo at marker** is the background subtracted average fluorescence of receptor at endosomes within the same ROI. Values at individual segmented endosomes within that ROI are averaged. (multiple lines, per ROI)

**Average per region norm Fluo at marker** is the background subtracted average fluorescence of receptor at endosomes normalized by the background subtracted average fluorescence of the receptor within the same ROI. Normalized values at individual segmented endosomes within that ROI are averaged. (multiple lines, per ROI)

**Total Av per region Fluo at marker** is the background subtracted average fluorescence of receptor at endosomes within the same ROI. This averaging is done by averaging all pixels for endosomes (compared to averaging individual endosomes) within the ROI.

**Total Av per region Norm Fluo at marker** is the background subtracted average fluorescence of receptor at endosomes normalized by the background subtracted average fluorescence of the receptor within the same ROI. This averaging is done by averaging all pixels for endosomes (compared to averaging individual endosomes) within the ROI.

**Per endosome Fluo at marker** is the background subtracted average fluorescence of receptor at endosomes across all ROIs. SEM is provided.

**Per endosome Norm at marker** is the background subtracted average fluorescence of receptor at endosomes across all ROIs, normalized by the background subtracted average fluorescence of the receptor in the corresponding ROI. SEM is provided.

<u>Fluorescence at individual endosomes is found in sheet 2</u>. First column will be the region ID, first line is average with sem below (if missing endosomes in one region this line gets buggy, sorry I did not fix this…but basically it is the average of all endosome values across all regions and associated sem). **This is the dataset that was used for further analysis in this manuscript with PoolManonBin.**

<u>Normalized Fluorescence at individual endosomes is found in sheet 3</u>. is the background subtracted average fluorescence of receptor at every endosomes across all ROIs, normalized by the background subtracted average fluorescence of the receptor in the corresponding ROI.

<u>Sheet 4</u> contains all parameters used for the quantification, including thresholds and polygon coordinates.

Program also generates a .tif 8 bit movie “name of your synapse movie_mask.tif” that contains the generated segmentation as follow:
polygon value = 1;

Receptor mask value = 2;
Endosome mask value = 3;

In the current manuscript, data were binned by time intervals, normalized and pooled. To do so, Excel files generated by the "Manon" script are then sorted into separate folders per condition. Set the current directory in the directory that contains these folders and enter "**PoolManonBin**" in the command window.
Code goes directly into "sheet 2" of each excel file, for each region it will obtain the average value from all detected structures of the first 6 frames. All values for this region are normalized by this average, and this is repeated for all regions. Normalized values are then averaged per bins frame 0-5, 6-10, 11-15, 16-20, 21-25, 26-30, 31-35 (time lapse were 35 minutes in these experiments). This is repeated across Excel files present in the folder, and code will output a "NameofFolderManon_PoolBin.xlsx" file that contains in the first column the name of the excel file pooled and the binned values for each file, together with an average and sem.

#### Analysis of SEP unquenching:

The 4 images per acquisition were assembled into a multi .tif stack. In a folder that contains this stack together with the first image, run "**ManuAllign**" in the command window. Code asks for the reference image (first image in our case), then for the stack to align (the 4 images stack). Reference images appear in green on the user interface, red is the stack to align. You will find controls to set the contrast and the frame, as well X and Y adjust that allow you to move each frame of the stack to it is corrected for drift compared to the reference image. When satisfied with the drift correction click on "export" and script will generate a .tif file "originalstackname_alligned".

Code will erase pixels (value set to 0) as you move the image to keep dimensions consistent. These areas were excluded from quantification by staying away from the edges.

To do the linescan analysis on the stack, we used a 4 images stack of the original syp-mCh image (duplicated 4 times) and the aligned stack. Set the current directory in a folder that contains these two files, enter "**Linescan2**" in the command window. Code asks for the stack to quantify first (in green) then for the synapse image (in red) that will appear color coded as indicated on the user interface.

This interface has been my "do everything" interface and contains many buttons, most of which will be buggy when transferred from one code to another, therefore I will not develop here what these buttons do and will focus on the ones that are relevant to this study.

You will find controls to set the contrast and frame as usual for this interface. Click on "axon" button to start manually drawing a linescan using the cross to select points with left click, to stop drawing use the right click for the last point. You might want to draw other lines, just repeat using the "axon" button. Do not change frame, and do not try to interact with the interface when drawing. When done selecting new lines, click on "Quantax" button, code will ask for the width of the linescan (3 in the current study), then select a rectangular background region and double click on it to validate. When done the code will display all kind of boxes around the lines that were visual controls for the linescan operation (no option to set the width of linescan with MATLAB, we had to rotate a rectangular matrix etc.). Script generates an Excel file "line_nameofthefile.xlsx" that contains the average fluorescence value of the linescan for each frame of the stack. Here is how it is calculated:

For each pixel along the line, an average is done along the width of the linescan. Then all linescans values are averaged by the total length of all the different linescans.

#### Analysis of SEP surface fluorescence:

In a current directory that contains your marker multi-tif file (synapse) and your surface fluorescence multi-tif file of the same number of images (SEP channel), enter "**CircleQuant2**" in the command window. Code asks for the SEP stack first (in green) then for the synapse image (in red) that will appear color coded as indicated on the user interface. To select synapses click on "Pick …", script will ask to select the background region, select synapses with left click, use right click when you are done selecting all synapses on the image. Move on to the next frame using the bottom left slider, repeat operation for each frame by clicking on "Pick …". When the whole stack is reviewed, click on "Quant …", code asks for the diameter of the circle in which to quantify the fluorescence (default is 3, what we used in the current study). An Excel file "nameofgreenchannel_CircleQuant2.xlsx" is generated. Every line of the file displays name of the green stack, frame, synapse number, X coordinate, Y coordinate, average fluorescence in the green channel, average fluorescence in the red channel.

This interface has been my "do everything" interface and contains many buttons, most of which will be buggy when transferred from one code to another. I you need to save the quantification before you are done reviewing the whole stack you can save the data and pick up where you left by going directly to the frame where you stopped, but make sure to change the name of the first saved .xlsx file or it will be edited and the quantification lost.

To estimate confidence intervals we used the script "**PermutToxls**". Variables for each measurement (list of fluorescence values) need to be saved beforehand as a .mat file. When running the code, enter the number of iterations N (50,000 in this study) and the name of the two variables you want to compare. The script generate N random bootstraps from each sample distribution and calculate the mean, and outputs the 2.5% (CI low) and 97.5% (CI high) (95% confidence interval values) in a excel file. The code also generate random permutation statistics on a N permutations, code retrieved from Laurens R Krol (2022). Permutation Test (https://github.com/lrkrol/permutationTest), GitHub. Retrieved May 24, 2022 but not used in the present study.

#### Analysis of SEP fluorescence bursts:

Movies are reviewed manually in a directory containing your synapse image and the SEP channel for the apparition of fluorescence bursts in the green channel using the "**CircleQuant2**" script as described above. Using the "Pick" button (lower one), events are selected for the frame where the burst of fluorescence appear. When done reviewing the stack, click on "save" and program will output a "nameofthegreenstack_annotate.txt" file that contains the events coordinate (event ID, frame, X coordinate, Y coordinate). To review the events and quantify the fluorescence, enter "**FT1cTIF**" in the command window. This script is adapted from DOI: https://doi.org/10.1523/JNEUROSCI.0799-14.2014. Code will ask to input parameters and displays a user interface the allows to browse selected events and displays quantifications. Click "next" to review all events, click "writeXLS" to export the quantification in a "nameofthegreenchannel_data.xls" file. The quantification is to be found in the "green sheet", are displayed the quantification parameters, the integrated fluorescence intensity in a "radius" defined circle around the selected event with the baseline (average fluorescence in the same area for the "baseline" frames before). Area (for average fluorescence) is found in the excel sheet. We used an unreasonably high "threshold parameter" (50) in this study to maintain the size/location of the fluorescence quantification. We obtained the average fluorescence by dividing the integrated intensity by the corresponding area value.
